# A Saturated Aminotriester Lipid Enhances Amonafide Activity in Human Colorectal Cancer Cells

**DOI:** 10.64898/2026.09.29.755247

**Authors:** Evy Hsen, Elizabeth Yu, Chancie Chou, Arin Kasliwal, Lucas Quan, Anna Gribok, Malavika Venkatesh, Nicholas Huang, Jayen Joshi, Adi Deodhar, Tiffany Zhang, Alina Liu, Alyssa Yee-Xin Chia, Yeonho Noh, Sameeha Mujtaba, Joseph E. Pazzi, Edward Njoo

**Affiliations:** Department of Chemistry, Biochemistry & Physical Science Aspiring Scholars Directed Research Program, Fremont, CA; Formulations, reThink64 Bionetworks PBC, Fremont, CA

**Keywords:** amonafide, aminotriester lipid, excipient, small-molecule delivery

## Abstract

The delivery of small-molecule therapeutics across cellular membranes remains a fundamental challenge in drug development, particularly for compounds whose physicochemical properties limit intracellular bioavailability. Inspired by the low toxicity, biodegradability, and synthetic accessibility of triethanolamine-derived lipids, we investigated a series of C18 aminotriester lipids as excipients for the codelivery of anticancer therapeutics. Four structurally related triester lipids bearing saturated, *cis*-monounsaturated, *trans*-monounsaturated, and diunsaturated C18 tails were synthesized through straightforward esterification reactions with triethanolamine. These lipids were evaluated for their ability to enhance the antiproliferative activity of several anticancer agents, including paclitaxel, doxorubicin, amonafide, homoharringtonine, camptothecin, and SN-38. Across multiple drug classes, the aminotriester lipids exhibited minimal intrinsic cytotoxicity and generally produced limited enhancement of drug potency. However, the fully saturated *tris*-stearate aminotriester lipid selectively enhanced the antiproliferative activity of amonafide in HCT-116 colorectal cancer cells, particularly at lower concentrations. In contrast, unsaturated lipid analogues displayed negligible enhancement effects, and the activity of the *tris*-stearate lipid was substantially attenuated in A549 lung adenocarcinoma cells, weakly observed in HT-29 colorectal cancer cells, and absent in CT26 murine colorectal cancer cells. Lactate dehydrogenase release assays indicated that these effects were not associated with membrane lysis or nonspecific cytotoxicity. Further, fluorescence microscopy time course experiments suggest improved membrane trafficking of amonafide when codelivered stoichiometrically with fully saturated C18 aminotriester lipids. Collectively, these findings demonstrate that saturated triethanolamine-based aminotriester lipids may function as selective and biocompatible codelivery excipients for certain small-molecule therapeutics. The observed dependence on both lipid saturation state and cellular context suggests that membrane organization and lipid packing may play important roles in mediating these effects. These results establish a foundation for further mechanistic investigations into aminotriester lipid-assisted small-molecule delivery systems.

## 1. Introduction

The delivery of therapeutic cargoes [1], including mRNA [2–4], small molecules, aptamers [5], and other oligonucleotide-based therapeutics [3,6,7], most fundamentally requires the enablement of a mono- or polyanionic payload to be delivered across a lipophilic membrane bilayer [8]. The delivery platforms may complex with anionic targets to form nanoparticles, liposomes, and other supramolecular structures, and these formulations have been previously described to be highly efficacious in mRNA vaccination [9], gene delivery, siRNA delivery [10,11], and small molecular drug delivery [12]. Among small molecules, which encompass a broad range of lipophilicity profiles, successful membrane delivery is contingent on a careful balance between lipophilicity-driven membrane permeability and hydrophilicity-driven solubility in both *intra-* and *extra*-cellular aqueous medium. Previously, many have investigated lipophilic excipients to enhance cytosolic bioavailability, and hence potency, of small molecule therapeutic agents. These lipophilic agents typically augment small molecule cellular uptake by acting as emulsifiers, cosolvents, solubilizers, or permeability enhancers [13]. While these strategies have demonstrated clinical success, a careful balance must be met in avoiding systemic or off-target toxicity imbued by the lipid delivery agent itself [14,15]. Triacylglycerols, as an example, are incredibly bio-compatible, with limited toxicity when used as a lipid-emulsifier excipient [16].

Owing to their exceptionally low inherent toxicity (LD_50_ > 700 mg/kg) [17–45], biodegradability, and simplicity of chemical preparation on scale, aminotriester lipids based on triethanolamine have previously been applied as surfactants and additives in a variety of chemical, industrial, and biological contexts, including as fabric softeners, toiletry additives, cosmetic products, and mRNA delivery agents [17–45]. In a recent example of this, Li and coworkers demonstrated that triethanolamine triesters can be prepared biocatalytically and are effective additives to lipid nanostructures for the delivery of mRNA payloads relevant to SARS-CoV-2 vaccination efforts [46]. Importantly, these and other studies demonstrated the unique capacity of C18-lipids in engaging biological membranes for enhanced drug delivery [47]. The diversity of naturally occurring unsaturated forms of these C18-lipids, which possess unique physicochemical properties [48], has further been demonstrated to play an important role in the efficacy of C18-lipid based delivery platforms [47]. For example, zwitterionic lipids such as 1,2-distearoyl-sn-glycero-3-phosphocholine (DSPC) and 1,2-dioleoyl-sn-glycero-3-phosphocholine (DOPC) [49], which are commonly used in formulations involving nucleic acid delivery, differ only in the presence of a *cis-*unsaturated olefin in the di-oleyl ester variant, but have vastly different biological properties [50].

Inspired by this, we sought to evaluate whether these triethanolamine-based triester lipids may enhance the *in vitro* activity of small molecule anticancer therapeutics. In light of the natural abundance and availability of unsaturated and saturated C18-lipids, we prepared a *tris-*stearate aminotriester lipid, which has no unsaturation along its C18 hydrocarbon tail, alongside a *tris-* oleate aminotriester and *tris-*linoleate aminotriester lipid, to investigate the role of adding a second *cis-*double bond to the alkyl tail. Further, to evaluate whether *cis-* and *trans-*unsaturated aminotriester lipids may possess differences in biological and biophysical behavior, we prepared the corresponding *tris-*elaidate aminotriester. In this study, this systematic series of C18 aminotriester lipids was probed for activity in improving the potency of paclitaxel (Taxol™) [51], an FDA-approved tubulin inhibiting antimitotic agent; doxorubicin [52], a DNA-intercalating topoisomerase II inhibitor; amonafide, a preclinically investigated topoisomerase II inhibitor [53]; homoharringtonine, a protein synthesis inhibitor [54]; camptothecin, a naturally-derived topoisomerase I inhibitor [55]; and SN-38, a potent derivative of camptothecin also functioning through Topoisomerase I inhibition [56]. We found that a *tris-*stearate aminotriester lipid enhanced potency of amonafide in HCT-116 cells, while the unsaturated lipid counterparts exhibited minimal potency enhancement across these compounds. Leveraging the intrinsic fluorescence of amonafide, we utilized fluorescence microscopy to track relative intracellular uptake of amonafide. Further, we found that this effect is specific to amonafide, as compounds with higher and lower logP identities did not benefit from *tris*-stearate codelivery. Finally, we show that the enhanced potency of amonafide imbued by the presence of the *tris*-stearate lipid is significantly attenuated in A549 lung cancer cells and had minimal effect in CT26 murine colorectal cancer cells. This suggests that the improved delivery of amonafide mediated by a *tris*-stearate lipid derivative of triethanolamine is payload- and cell-line-dependent.

## 2. Materials and Methods

### 2.1 Cell Culture

HCT-116, HT-29, CT26, and A549 cells were cultured in T25 and T75 flasks. The cultures were kept at 37°C in a humidified incubator (5.0% CO_2_) and maintained by splitting cells at a ratio of 1:3 to 1:6 based on confluence and cultured up to passage 20. HCT-116 and HT-29 human colorectal cancer cell lines were cultured in McCoy’s 5A Medium (Tribioscience) supplemented with 10% v/v fetal bovine serum (FBS, Gibco) and 1% v/v 100x penicillin-streptomycin (Tribioscience). CT26, a murine colorectal carcinoma cell line, was cultured in RPMI-1640 Medium (Tribioscience) supplemented with 10% v/v fetal bovine serum (FBS, Gibco) and 1% v/v 100x penicillin-streptomycin (Tribioscience). A549, a human non-small cell lung carcinoma cell line was maintained in High Glucose Dulbecco’s Modified Eagle Medium (DMEM) (Hygia Reagents) supplemented with 10% fetal bovine serum (Gibco) and 1% penicillin-streptomycin (Tribioscience). HCT-116 cells were purchased from European Collection of Authenticated Cell Cultures (ECACC), Millipore Sigma Cat. # 91091005-1. HT-29 cells were purchased from European Collection of Authenticated Cell Cultures (ECACC), Millipore Sigma Cat. #91072201. A549 cells were a generous gift from Dr. Feng Wang-Johanning and Dr. Gary Johanning (SunnyBay Biotech, Fremont, CA) and were originally purchased from ATCC. CT26 cells were a generous gift from Dr. Zhong Wang (BJ Biosciences) and were originally purchased from ATCC.

### 2.2 MTT Assay

Cell viability was determined by 3-(4,5-dimethylthiazol-2-yl)-2,5-diphenyltetrazolium bromide (MTT) assays through literature reported protocols [57,58]. HCT-116 and HT-29 cells were seeded at 70% confluency in McCoy’s 5A Medium and supplemented with 10% v/v fetal bovine serum (FBS, Gibco) and 1% v/v 100x penicillin-streptomycin (Tribioscience). A549 and CT26 cells were seeded at 70% confluency in High Glucose Dulbecco’s Modified Eagle Medium (DMEM) and RPMI-1640 respectively, supplemented with 10% v/v fetal bovine serum (FBS, Gibco) and 1% v/v 100x penicillin-streptomycin (Tribioscience). Following the 24-hour incubation period, a drug medium solution was prepared by pre-formulating our triethanolamine triester lipids with select chemotherapeutics. Equivalent volumes of drug and lipid solutions were mixed in a v-bottom 96-well plate, which were added to media in a deep-well plate. 100 µL of the drugged media was then added to each well to reach desired concentrations of 25 µM, 5 µM, 2.5 µM, 500 nM, 250 nM, 50 nM, 25 nM, and 5 nM (0.5% v/v dimethyl sulfoxide (DMSO)). A negative control of 0.5% v/v DMSO was also included. The plates were then incubated at 37°C (5.0% CO2) for 96 hours, after which 10 μL of a fresh solution of 3-(4,5-dimethylthiazol-2-yl)-2,5-diphenyltetrazolium bromide (MTT) (AK Scientific) in 1x phosphate-buffered saline (PBS) (5 mg/mL) was added to all wells. After mixing, the plates were allowed to incubate for 1-3 hours. The cell media was then aspirated, and 100 μL of DMSO was added to each well and mixed until all formazan crystals were fully solubilized. Absorbance was measured with a Molecular Devices SPECTRAmax 250 Microplate Spectrophotometer at 570 nm. Cell viability (%) was calculated and normalized against the negative 0.5% v/v DMSO control and plotted against drug concentration. IC_50_ values were then determined using GraphPad Prism 10.4.1 with an inhibition regression analysis.

### 2.3 LDH Assay

Extracellular LDH activity, released due to a loss of cell membrane integrity, was measured using a previously reported Cold Spring Harbor Protocol. HT-29 cells were seeded at 70% confluency in McCoy’s 5A Medium and supplemented with 10% v/v fetal bovine serum (FBS, Gibco) and 1% v/v 100x penicillin-streptomycin (Tribioscience) [59]. Following the 24-hour incubation period, a drug medium solution was prepared by pre-formulating our triethanolamine triester lipids with select chemotherapeutics. Equivalent volumes of drug and lipid solutions were mixed in a v-bottom 96-well plate, which were added to media in a deep-well plate. 100 µL of the drugged media was then added to each well to reach desired concentrations of 25 µM, 5 µM, 2.5 µM, 500 nM, 250 nM, 50 nM, 25 nM, and 5 nM (0.5% v/v DMSO). A negative control of 0.5% v/v DMSO was also included. The plates were then incubated at 37°C (5.0% CO2) for 96 hours. At 96 hours, to a negative control of cells drugged with DMSO was added 10 μL of lysis solution (9% v/v Triton X-100) for 5 minutes to act as a positive control for maximum LDH release. 50 µL of supernatant cell media from each treatment was then moved to a non-tissue culture treated 96-well plate and allowed to react with 50 µL of the LDH substrate solution (L-(+)-lactic acid (0.054 M), β-NAD+ (1.30 mM), 1-Methoxy-5-methylphenazinium methyl sulfate (0.28 mM), and 2-p-iodophenyl-3-p-nitrophenyl tetrazolium chloride (INT) solution (0.66 mM) dissolved in 0.2 M Tris-HCl buffer (pH 8.2), for 45 min at 37°C (5.0% CO2) protected from light. Absorbance was measured with a Molecular Devices SpectraMax® Plus 384 Microplate Reader at 490 nm.

### 2.4 Fluorescence microscopy

HCT-116 and CT26 cell lines were cultured and utilized to evaluate the fluorescence localization of amonafide in the presence of four lipid formulations. Stock solutions consisted of 10 mM amonafide dissolved in DMSO, four lipid formulations, *tris*-stearate, tris-oleate, *tris*-linoleate, *tris*-elaidate prepared in PBS, and 1 mg/mL mixture of 1,2-Dioleoyl-sn-glycero-3-phosphocholine (DOPC): 1,2-Dioleoyl-sn-glycero-3-phosphoethanolamine-N-(lissamine rhodamine B sulfonyl) (RhodamineDOPE) (95.28:4.72mol%). For each experimental condition, 10 μL of amonafide solution and 20 μL of the designated lipid formulation were added to individual wells. Experiments were conducted three times for each lipid formulation and cell type. Each trial consisted of four treatment groups: (1) amonafide alone, (2) amonafide followed by lipid addition after a 5 min period, (3) lipid alone, and (4) lipid followed by amonafide addition after a 5 min period. After all the compounds were added, all treatment groups were incubated for an additional 5 minutes before proceeding with the assay. Thus, each lipid formulation was evaluated using 12 wells (4 treatment groups 3 replicates). Across all four lipid formulations, a total of 48 wells were analyzed per cell line, resulting in 96 experimental wells across both HCT-116 and CT26 cell types. Immediately prior to imaging, approximately 150 μL of medium was carefully removed from each well, leaving a minimal volume sufficient to prevent cells from drying out during imaging. Fluorescence microscopy was performed using a 10x objective lens [60]. For each well, three images were acquired from the same field of view. First, a brightfield image was captured using the white filter taken using halogen. Then, fluorescence images are taken using the green filter set followed by the red filter set, ensuring that all images were collected from identical imaging locations. Fluorescence was used exclusively during the green and red channel images. A total of three images (brightfield, green fluorescence, and red fluorescence) were taken for every experimental well.

### 2.5 Critical Micelle Concentration Measurements

Apparent critical micelle concentration measurements were performed using a contact angle assay [61]. To begin, 20 μl of a 50 mM compound stock in DMSO was evaporated to dryness in a SpeedVac (Savant SC100) for approximately 2 hours, then reconstituted with 20 μl of 0.5% DMSO in water. Samples were bath sonicated (Thermo Fisher) for 10 min and briefly vortexed. Ten-fold serial dilutions from 50 mM to 5 nM were then prepared in a Falcon 96-well clear round-bottom (U bottom) polystyrene microplate by transferring 2 μL of compound solution into 18 μL of 0.5% DMSO in water, with four replicate wells prepared per concentration. For each concentration, 2 μL droplets were taken from each replicate well and placed onto a Teflon surface positioned in front of a blue background. A 0.5% DMSO control was included. Droplets were imaged at 5x zoom, and contact angles were measured in ImageJ at 180° minus θ_E_. The apparent CMC was determined from the concentration-dependent change in contact angle across the dilution series.

### 2.6 Dynamic Light Scattering data

Z-Average hydrodynamic radii were measured with Protein Solutions DynaPro-99-E-50 System Dynamic Light Scattering Module. Stock solutions with 10 mM of aminotriester lipids in DMSO were prepared. Then, half log serial dilutions of amonafide in DMSO from 10 mM to 316 nM were prepared. Finally, the test solutions were generated by mixing 4 μL of the aminotriester lipid stock, 4 μL of the amonafide dilution of the target concentration, and 1592 μL of DI water, yielding a final solvent composition of 0.5% DMSO. Additionally, mixtures of 0.25% v/v aminotriester lipid stock in DI water were formulated as a control. Each solution was tip-sonicated in bursts for four minutes prior to measurement.

## 3. Results

### 3.1 Chemical synthesis

In order to evaluate the effect of lipid tail saturation in the function of triester lipids, we prepared a series of four triester lipids with varying numbers of alkenes. This was accomplished with standard carbodiimide-mediated esterification reactions to triethanolamine [62], affording the corresponding saturated, monounsaturated, and polyunsaturated triester lipids (**Figure 1**). Full characterization, including ^1^H and ^13^C NMR spectra, is included in the electronic Supporting Information document.

**Figure 1.**
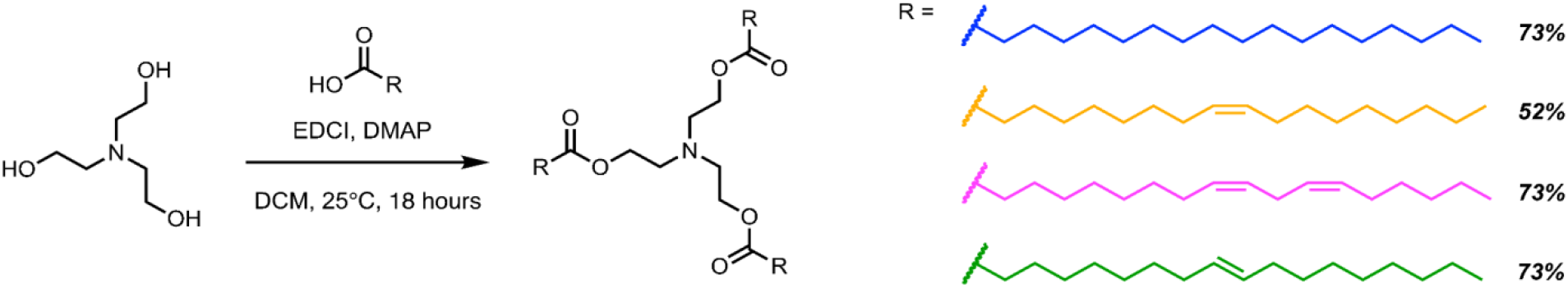
Chemical synthesis of C18 triester triethanolamine lipids. Tripod lipids bearing *tris*-stearate, *tris-*oleate, *tris-*linoleate, and *tris-*elaidate C18 alkyl chains were prepared to interrogate the effect of mono- and poly-unsaturation as well as *cis-* and *trans-*unsaturated C18 lipids.

### 3.2 Evaluating antiproliferative activity in co-delivery assays

To determine whether any improvement in biological activity of our codelivery systems was attributed to the inherent toxicity of the triester lipids or the enhancement of drug delivery, we first evaluated the cytotoxic effects of our triester lipids through a preliminary cell viability assay against a panel of cancer cell lines including HCT-116, HT-29, and A549 cells. In this assay, our lipids demonstrated a lack of cytotoxicity, suggesting their potential as structural components of larger drug delivery systems. Furthermore, we evaluated the stearate lipid in the HCT-116 colorectal cancer cell line and found that the lipid alone did reduce cell viability. We subsequently co-delivered our lipids in 3 additional cell lines, comparing the IC_50_ with and without the addition of our lipids. Upon analysis, we discovered that amonafide, when co-delivered with *tris*-stearate, has a statistically significant improved potency compared to its administration alone. With this lead, we proceeded to evaluate the ability of our stearate triester lipid to improve amonafide activity across four cell lines. Interestingly, there appears to be an amplification in our stearate lipid’s effect at lower concentrations (2.5 μM to 5.5 nM) and a suppression at higher concentrations (5 μM to 25 μM) in both the HCT-116 and A549 cell lines, suggesting that the lipid-amonafide ratio has significant influence on the observed effect. In the HT-29 cell line, the enhancement of amonafide potency was less pronounced, and in the CT26 cell line, there was a more attenuated effect, which could be due to difference in membrane composition and sensitivity to permeabilizing agents.

### 3.3 Cell Viability Assay

To quantify and identify a potential enhancement in potency of small molecule anticancer therapeutics through co-delivery with the triester lipids, we measured 96-hour IC_50_ values for six small molecule toxins with distinct mechanisms of action: doxorubicin, paclitaxel, camptothecin, SN-38, amonafide, and homoharringtonine. Each compound was evaluated for anticancer activity when administered alone or with one of the four triester lipids in HCT-116, HT-29 (human colorectal carcinoma), and A549 (human lung adenocarcinoma) cell lines (**Figure 2)**. As shown in **Figure 2**, the majority of drug-lipid pairs shifted IC_50_ by less than threefold relative to the drug alone. Amonafide co-delivered with *tris*-stearate was an exception, as its IC_50_ values decreased from 2547 to 26.9 nM in HCT-116, from 1089 to 3.14 nM in HT-29, and from 633 to 3.36 nM in A549 cell lines.

**Figure 2:**
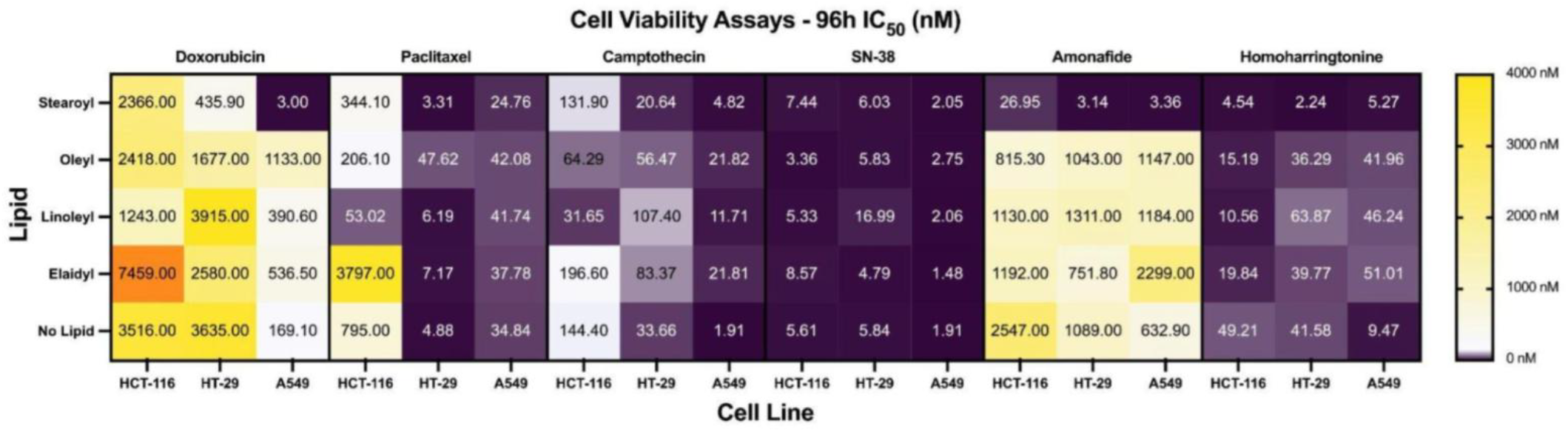
Antiproliferative activity of six anticancer drugs in HCT-116, HT-29, and A549 cells when co-delivered with tripod lipids. IC_50_ heatmap of anticancer therapeutics co-delivered with C18 triester triethanolamine lipid. Doxorubicin, paclitaxel, camptothecin, SN-38, amonafide, and homoharringtonine were co-delivered with all four lipids and compared against a negative control in HCT-116, HT-29, and A549 cell lines.

Expanding on the apparent special nature of the amonafide and *tris*-stearate co-delivery, we further examined the effect of *tris*-stearate on the inhibition of amonafide across the prior cell lines, with the additional inclusion of CT26 (murine colon carcinoma) (**Figure 3**). As shown in **Figure 3B**, codelivery with *tris*-stearate significantly reduced amonafide-induced viability relative to amonafide alone at nearly every concentration tested in HCT-116, HT-29, and A549, consistent with the substantial IC_50_ shifts demonstrated in the heatmap from **Figure 3A**. Specifically, in HCT-116, the IC_50_ decreased from 27.6 µM to 71.7 nM, a 385-fold enhancement in potency and by far the largest observed among the four cell lines. Moreover, these differences in viability were statistically significant across the sub-micromolar ranges. A similar pattern was observed for HT-29, with enhanced potency across the low to moderate concentration ranges. However, the corresponding shift in IC_50_, from 28.1 µM to 9.73 µM, was comparatively modest, with only a 2.9-fold change. In A549 cells, the codelivery with *tris*-stearate also significantly reduced viability at sub-micromolar concentrations (**Figure 3B**). This corresponded to an approximately 111-fold decrease in IC_50_, from 1.45 µM to 13.1 nM (**Figure 3A**). Clearly, the delivery in the presence of *tris*-stearate improved the potency, although at a smaller magnitude than observed in HCT-116. CT26 cells were the least responsive to codelivery, as seen by the only 1.4-fold shift in IC_50_ (**Figure 3A**), as well as the minor shifts in viability recorded across all concentrations (**Figure 3B**). Taken together, these results show a clear hierarchy in the *tris*-stearate mediation of enhanced amonafide activity, where the effect is most pronounced in HCT-116, substantial but modest in A549 and HT-29, and largely absent in CT26. This supports a cell line dependent potency enhancement.

**Figure 3:**
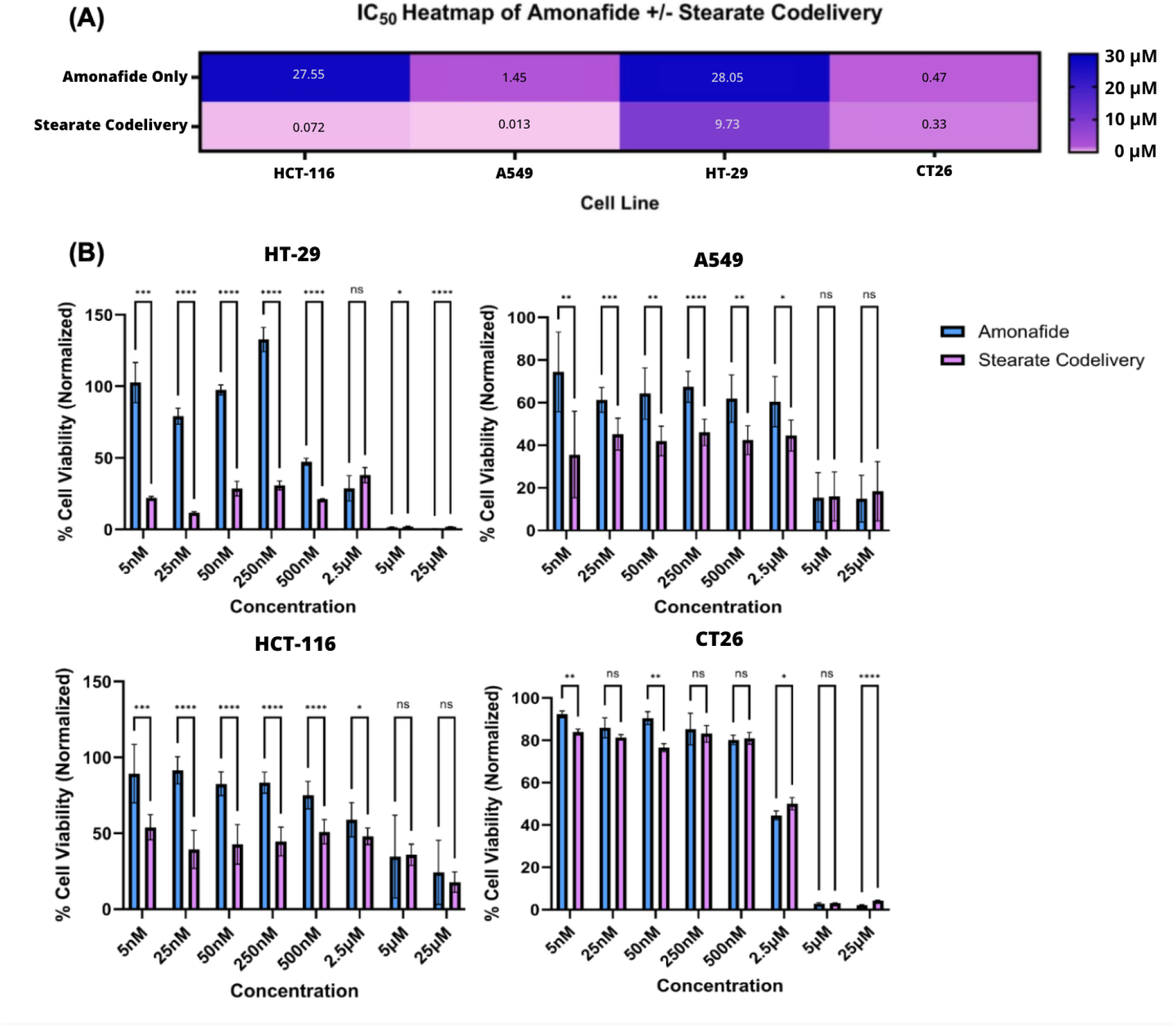
Cell viability in the presence and absence of codelivery with *tris*-stearate triethanolamine lipid. **A**. Having identified a unique trend of improved potency of amonafide delivered with *tris-*stearate tripod lipid, this is a direct comparison of amonafide delivered with and without the *tris*-stearate lipid in HT29, A549, HCT-116, CT26 cell lines. **B. HT**-29 (n = 8), A549 (n = 8), HCT-116 (n = 8), and CT26 (n = 4) were incubated with amonafide with and without codelivery of tris-stearate triethanolamine lipid for 96 hours prior to endpoint analysis. ns (nonsignificant), * < 0.05, ** < 0.01, *** < 0.001, **** < 0.0001.

LDH assays demonstrate that the codelivery of tripod lipids with these six selected anticancer compounds do not elicit significant differences in necrotic cell death, with similar trends of increasing LDH activity at higher concentrations of amonafide regardless of codelivery with each aminotriester lipid (**Figure 4**). This suggests that the improved activity of amonafide in codelivery with the *tris-* stearate tripod lipid is not attributed to significant changes in membrane integrity.

**Figure 4:**
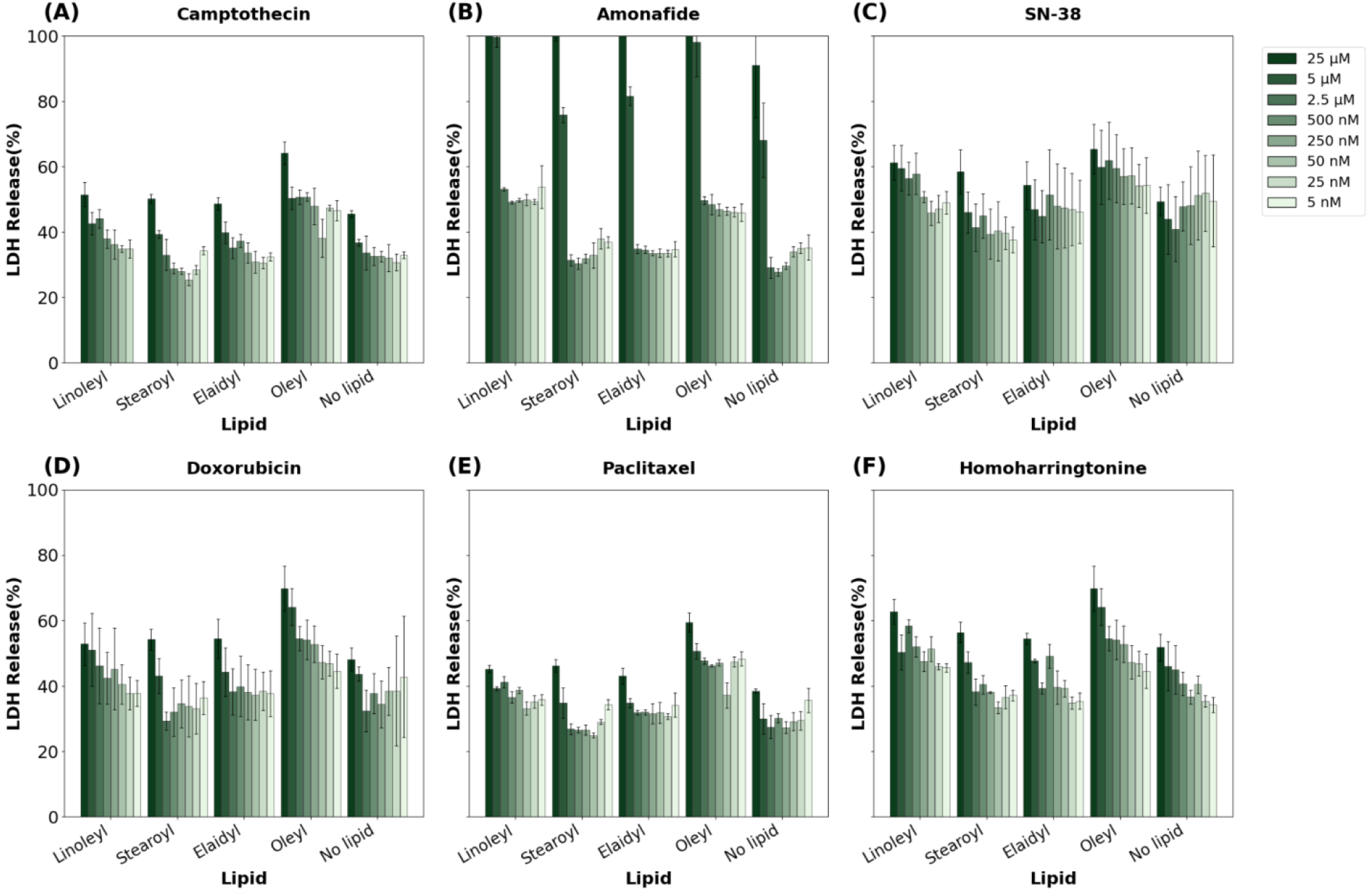
Cytotoxicity of camptothecin (A), amonafide (B), SN-38 (C), doxorubicin (D), paclitaxel (E), and homoharringtonine (F) observed through lactate dehydrogenase (LDH) assays in HT-29 human epithelial colorectal cancer cells. Data is represented as means ± SD compared with the negative control.

#### 3.4 Fluorescent-based Cell Uptake Assay

To further rationalize these biological results, we developed a biophysical characterization workflow that included fluorescence microscopy tracking of cellular uptake, critical micelle concentration measurements, and dynamic light scattering particle analyses. We first performed fluorescence microscopy to evaluate whether the selective potency enhancement of amonafide by *tris-*stearate was caused by intracellular drug accumulation. We quantified the mean fluorescence intensity per field using a custom image analysis routine developed in our previous work [63]. **Figure 5** shows brightfield and fluorescent images of HCT-116 and CT26 cells, which were treated under five conditions: i) untreated control, ii) amonafide alone, iii) a 5-minute amonafide incubation followed by aminotriester lipid, iv) lipid alone, and v) a 5-minute aminotriester lipid incubation followed by amonafide. Amonafide was visibly fluorescent in the green channel, allowing us to track intracellular drug delivery without requiring any additional staining procedures. The lipid analog mixtures were doped with a rhodamine-PE lipid and imaged in the red channel, which allowed for tracking of the lipid analogs to be confirmed independently of drug measurement. Brightfield images were also collected to match the fluorescence signal in the images to the location of the cells.

**Figure 5.**
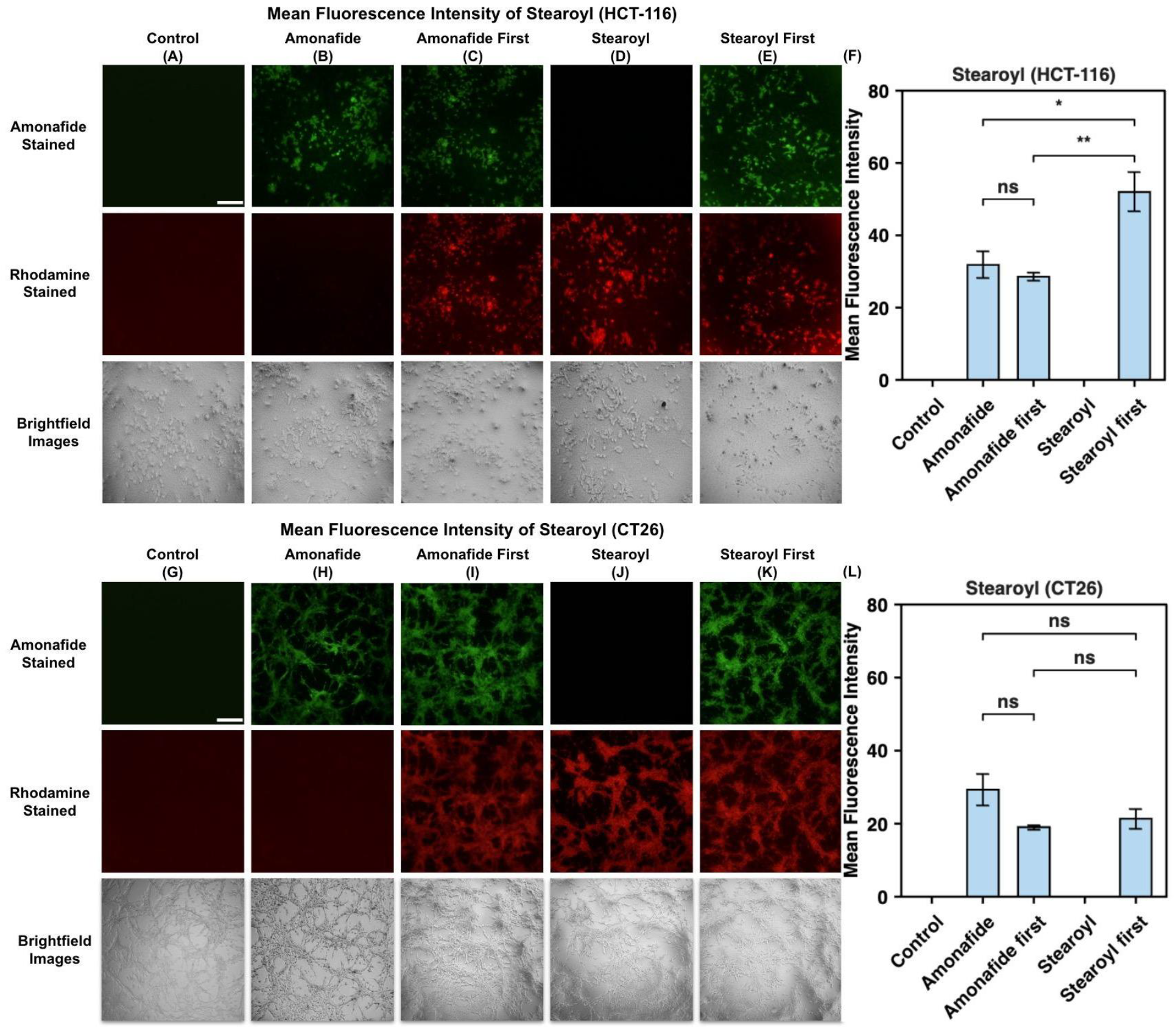
Representative fluorescence images and quantification of intracellular fluorescence signals in HCT-116 (A-F) and CT26 (G-L) cells. Columns correspond to treatment groups: untreated control (**A, G);** amonafide alone (**B, H)**; amonafide applied first, followed 5 min later by *tris*-stearate (**C, I**); *tris-*stearate alone **(D, J)**; and *tris*-stearate applied first, followed 5 min later by amonafide (**E, K**). The first row indicates the green channel, which signals arise from the fluorescence of amonafide and report intracellular drug. The red channel is indicated by the rhodamine-tagged lipid and reports lipid delivery. The brightfield images support the location of the fluorescent-dyed lipid cells in both green and red channels. Quantitative data were provided by bar graphs, which represent mean fluorescence intensity ± SEM (**F, L)**. Statistical significance was determined as indicated: ns (not significant), * < 0.05, ** < 0.01. Scale bars are 50 µm (**A, G)**.

Untreated cells produce no detectable signal in either fluorescent channel **(Figure 5A, 5G)**. Cells given the lipid-alone treatment produced a strong red signal, but no green signal above background, demonstrating that the lipid reaches the cell while not causing any artifacts in the amonafide channel **(Figure 5D, 5J)**. Thus, we attribute the signal observed in the green channel images to the presence of amonafide, which suggests that the differences in the intensities between the treatment groups are correlated to differences in intracellular drug concentrations.

In HCT-116 cells, the order of administration was only significant for *tris*-stearate but not for the unsaturated lipids (**Figure 5F, 5L**). Applying the *tris*-stearate first raised mean fluorescent intensity from 28.5 ± 1.1 (amonafide first) to 52.0 ± 5.4 (lipid first), a 1.8-fold increase. Neither sequence differed significantly from amonafide alone (31.9 ± 3.8, p = 0.43 and p = 0.11, respectively), suggesting that this effect was attributed solely to the application of *tris*-stearate before the drug. The three unsaturated lipids showed no significant sequence dependence in this cell line (*tris*-elaidate p = 0.33, *tris*-oleate p = 0.23, *tris*-linoleate p = 0.44; **SI 2-4**). The effect therefore appears strongest for the fully saturated lipid. In CT26, there is no statistically significant difference between the order of administration for the tripod lipids (*tris-*stearate p = 0.48, *tris*-elaidate p = 0.058, *tris*-oleate p = 0.083, *tris*-linoleate p = 0.88; SI **5-7**). This suggests that the significance seen in HCT-116 by *tris-*stearate is dependent on the cell line.

These results indicate that the measured fluorescence intensity originates from amonafide rather than the lipid carriers, and that of the four triester lipids, only *tris*-stearate produced a statistically significant change in intracellular amonafide signal, and only in HCT-116. Consequently, while the sequence of lipid-carrier administration can selectively enhance amonafide delivery in specific contexts, it does not universally modulate short-term intracellular localization across all lipid matrices within a five-minute incubation window.

### 3.5 Critical Micelle Concentration Assay

To determine whether the enhanced potency of amonafide in the presence of *tris*-stearate is caused by an alteration in lipid self-assembly, we measured the critical micelle concentration (CMC) of each triester lipid alone and then in the presence of amonafide. The apparent CMC was determined by calculating the contact angle across a ten-fold serial dilution series, where the initial drop of the curve represents the onset of self-assembly (**Figure 6)**. Due to the limited solubility of the lipids in 0.5% DMSO in water, concentrations above the tested range could not be evaluated.

**Figure 6.**
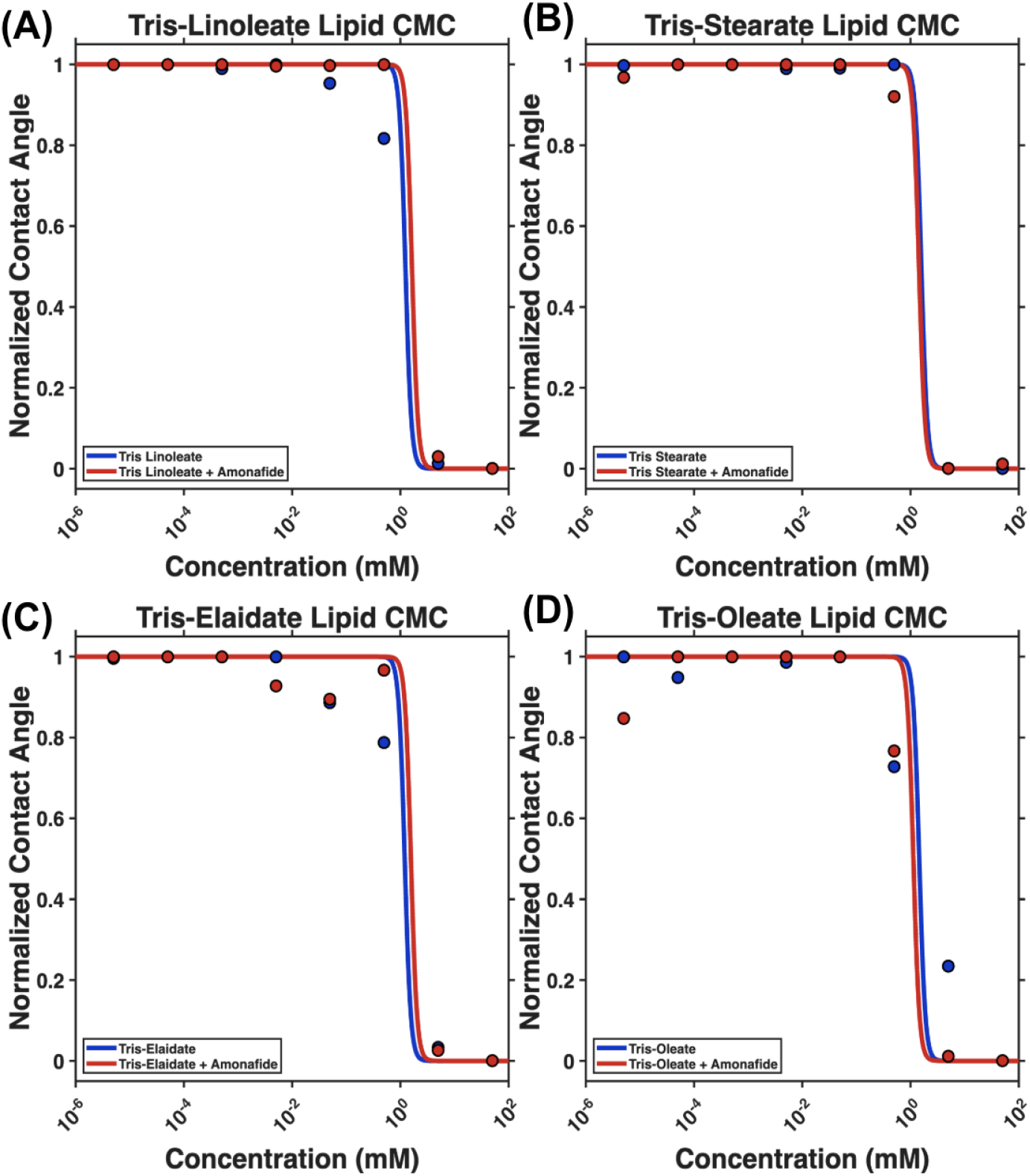
Critical micelle concentration of tripod lipids with and without amonafide. Normalized contact angle curves are presented for **(A)** *tris-*linoleate, **(B)** *tris-*stearate, **(C)** *tris-*elaidate, and **(D)** *tris-* oleate lipids with and without amonafide. Blue curves represent lipids alone, while red curves represent lipids with amonafide. Data points represent normalized final average contact angle values across the dilution series, and solid lines represent fitted sigmoidal CMC transitions.

The presence of amonafide did not significantly alter the transition points for any of the four lipids in this series, as shown in **Figure 6**. Importantly, *tris*-stearate (**Figure 6b)** showed no shift in apparent CMC when the amonafide and the lipid were combined, indicating that the presence of amonafide does not bear significant influence on the self-assembly propensity of the *tris*-stearate lipid. In connection to the earlier observed augmented biological effects, these data suggest that the increase in potency of *tris*-stearate is not caused by a shift in self-assembly into a supramolecular structure.

### 3.6 Dynamic Light Scattering Assay

While particle size has previously been demonstrated to be a key driver in cellular drug uptake [64], we aimed to understand how the specific particle dimensions and size regimes differ or compare across the four aminotriester lipids through dynamic light scattering experiments. We measured the Z-average hydrodynamic radius of the aminotriester lipid particles in the presence of varying concentrations of amonafide. As shown in **Figure 7**, the hydrodynamic radii of these particles were consistently similar to that of the self-assembled particles without amonafide, suggesting that amonafide was neither self-assembling with the aminotriester lipids nor affecting the self-assembly of the aminotriester lipids. This aligns with the CMC data and further supports our hypothesis that the enhanced potency of amonafide with *tris*-stearate observed in cells depends not on a perturbation in self-assembly but rather on a localized effect between drug and lipid.

**Figure 7.**
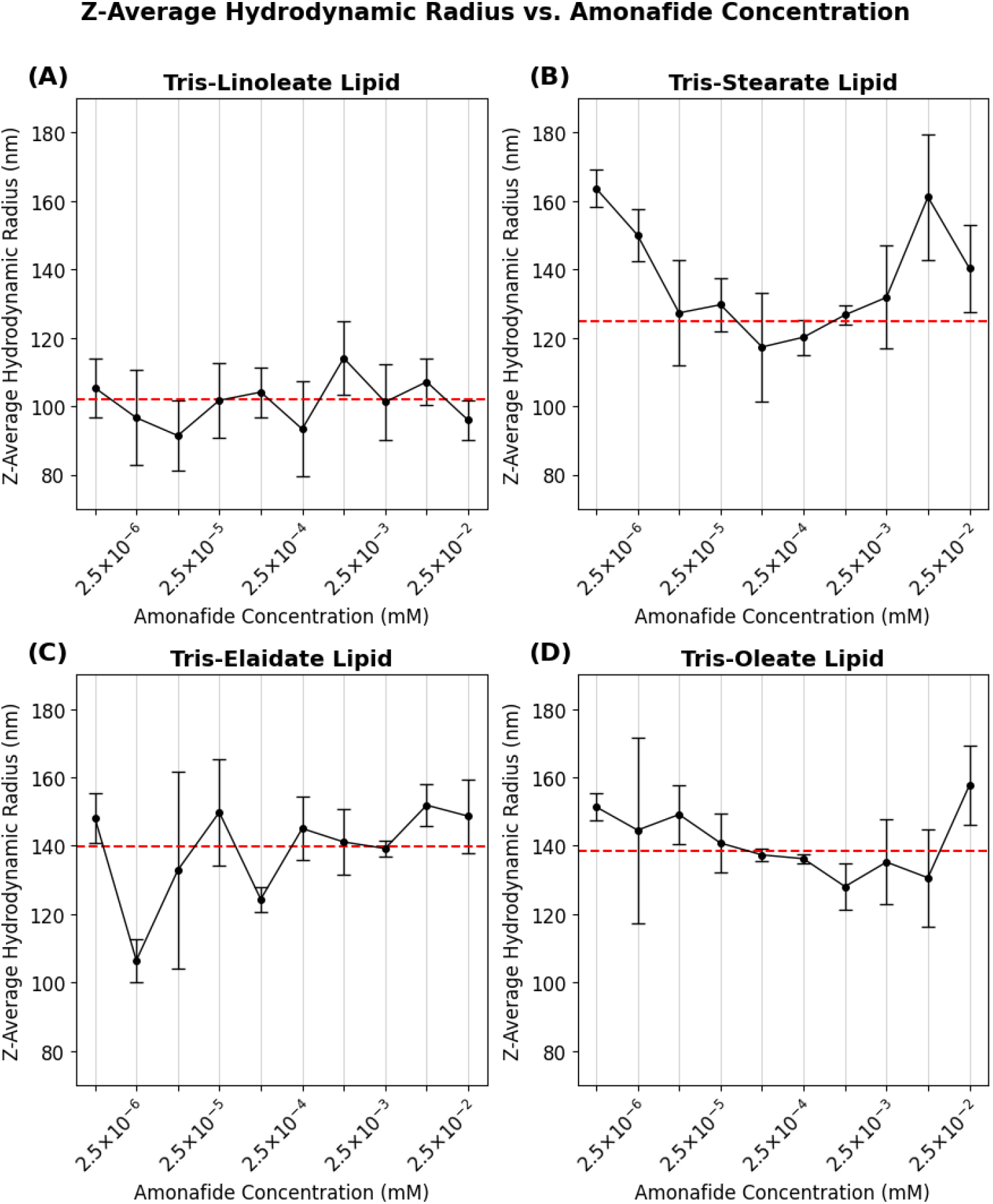
Z-average hydrodynamic radius measurements of aminotriester lipid particles across changing concentrations of amonafide. Aminotriester lipids measured are **(A)** *tris*-linoleate, **(B)** *tris*-stearate, (**C)** *tris*-elaidate, and **(D)** *tris*-oleate. Solid lines represent the Z-average hydrodynamic radius of the lipid particles with amonafide. Dotted lines represent the Z-average hydrodynamic radius of the lipid particles without amonafide present. Error bars represent the standard deviation.

After sonication, *tris*-stearate (**Figure 7A**) self-assembles into lipid particles with a Z-average hydrodynamic radius of ∼125 nm, *tris*-elaidate **(Figure 7D**) self-assembles into particles with a Z-average hydrodynamic radius of ∼140 nm, and *tris*-oleate (**Figure 7C)** self-assembles into particles with a Z-average hydrodynamic radius of ∼138 nm. As shown in **SI 8**, *tris*-stearate, *tris*-elaidate, and *tris*-oleate exhibit a wide particle size distribution. Conversely, as shown in **SI 8**, *tris*-linoleate has a particle size distribution with a higher population of particles with a smaller hydrodynamic radius. *Tris*-linoleate (**Figure 7B**) self-assembles into the smallest particles with a Z-average hydrodynamic radius of ∼102 nm, with a size distribution which is statistically significant (p = 6.53e-7) in comparison to the other three aminotriester lipids. Simultaneously, *tris*-linoleate expresses the most monodisperse distribution, while the other three aminotriester lipids exhibit much broader peaks. Due to the presence of its two *cis*-double bonds, *tris*-linoleate exhibits less lipid packing than monounsaturated and saturated lipids. This suggests that when sonicated, the lipids require less energy to separate and as a result form smaller initial lipid droplets before they self-assemble into lipid particles. Since the initial *tris*-linoleate lipid droplets are smaller, the lipid particles are more likely to have a smaller Z-average hydrodynamic radius.

## 4. Discussion & Conclusions

Inspired by the structural simplicity and biodegradability of ester-based lipid excipients, we investigated how C18 acyl-chain saturation in triethanolamine-derived aminotriester lipids influences their ability to enhance small-molecule anticancer activity. We evaluated these lipids in combination with a mechanistically diverse panel of anticancer agents, including paclitaxel, doxorubicin, amonafide, camptothecin, homoharringtonine, and SN-38. To systematically interrogate the contribution of lipid packing and unsaturation, we prepared a matched series comprising fully saturated *tris-*stearate, *cis*-monounsaturated *tris*-oleate, *trans*-monounsaturated *tris-*elaidate, and diunsaturated *tris-*linoleate aminotriesters.

Uniquely, the fully saturated tristearate ester of triethanolamine enhanced the potency of amonafide in HCT-116 human colorectal cancer cells, particularly at lower concentrations. This effect was significantly attenuated in other human cancer cells such as A549 lung adenocarcinoma cells and HT-29 and fully abolished in CT26 murine colorectal cancer cells, suggesting that there may be selectivity on certain cell lines, and that this phenomenon may be unique to the interaction of our *tris-*stearate aminotriester lipid with amonafide potency in human CRC cell lines. Expectedly, this and other triester lipids exhibited no inherent toxicity to cells, even at maximally soluble concentrations.

We found that the majority of these aminotriester lipid -small molecule codelivery examples demonstrated no significant enhancement of *in vitro* potency, suggesting that this effect may be limited to amonafide and compounds of similar biophysical features. Next, to interrogate the mechanism through which the *tris-*stearate lipid amplifies amonafide potency, we leveraged the intrinsic fluorescent properties of amonafide and conducted time-course fluorescence uptake experiments. The data obtained from these experiments suggest that, across both the HCT-116 and CT26 cell lines, while the mean fluorescence intensity indicates that changing the administration order or pairing the drug with a specific aminotriester lipid increases amonafide delivery, with marginal improvements depending entirely on the specific aminotriester lipid and cell line. This is consistent with the significant *in vitro* selectivity observed in the *tris*-stearate treatment. While other lipid formulations exhibited minor fluctuations in the mean fluorescent intensity under different sequences, the variations did not show statistical significance. Although the sequence of lipid administration can selectively enhance amonafide delivery in *tris*-stearate, it does not modulate intracellular localization across all aminotriester lipids within a 5-minute incubation window.

Furthermore, the Z-average hydrodynamic radii of aminotriester lipids and critical micelle concentration are not affected by the presence of amonafide, suggesting that the improved efficiency of amonafide in HCT-116 cell lines with *tris*-stearate is not due to the spontaneous discrete amonafide-lipid supramolecular structure assemblies. This does not preclude other potential mechanisms through which the *tris*-stearate lipid may be improving the anticancer potency through direct but non-toxic membrane delivery.

Collectively, these results demonstrate that fully saturated aminotriester C18 lipids have specific contexts in which they may improve the potency of small molecule anticancer agents. Further studies may shed light on the mechanistic basis for this unique activity in the delivery of the preclinical small molecule anticancer agent amonafide.

## Author Contributions

Conceptualization, J.P., E.N..; methodology, E.H., E.Y., C.C., A.K., L.Q., A.G., M.V., N.H., J.J., A.D., T.Z., A.L., A.Y.C., Y.N., J.P., E.N.; software, A.K., L.Q., M.V., N.H., J.J., A.D..; validation, E.H., E.Y., C.C., A.K., L.Q., A.G., M.V., N.H., J.J., A.D., T.Z., A.L., A.Y.C., Y.N., J.P., E.N.; formal analysis, E.H., E.Y., C.C., A.K., L.Q., A.G., M.V., N.H., J.J., A.D., T.Z., A.L., A.Y.C., Y.N., J.P., E.N.; investigation, E.H., E.Y., C.C., A.K., L.Q., A.G., M.V., N.H., J.J., A.D., T.Z., A.L., A.Y.C., Y.N., J.P., E.N.; resources, J.P., E.N.; data curation, E.H., E.Y., C.C., A.K., L.Q., A.G., M.V., N.H., J.J., A.D., T.Z., A.L., A.Y.C., Y.N., J.P., E.N.; writing— original draft preparation, E.H., E.Y., C.C., A.K., L.Q., A.G., M.V., N.H., J.J., A.D., T.Z., A.L., A.Y.C., Y.N., J.P., E.N.; writing—review and editing, E.H., C.C., A.K., L.Q., A.G., M.V., N.H., J.J., A.D., J.P., E.N.; visualization, E.H., E.Y., C.C., A.K., L.Q., A.G., M.V., N.H., J.J., A.D., J.P., E.N.; supervision, J.P., E.N.; project administration, J.P., E.N..; funding acquisition, J.P., E.N. All authors have read and agreed to the published version of the manuscript.

## Institutional Review Board Statement

Not applicable.

## Data Availability Statement

Experimental data and characterization information for compounds are contained in the Electronic Supporting Information (SI) document.

## Acknowledgments

The authors gratefully knowledge Dr. Stephen Lynch and the Stanford University NMR Facility for access to high field NMR spectroscopy. Additionally, the authors gratefully acknowledge Dr. Feng Wang-Johanning and Dr. Gary Johanning from SunnyBay Biotech for their generous donation of several cell lines used in this study.

## Abbreviations

The following abbreviations are used in this manuscript:

C18: Carbon-18
mRNA: Messenger ribonucleic acid
LD_50_: Median Lethal Dose
DSPC: 1,2-Distearoyl-sn-glycero-3-phosphocholine
DOPC: 1,2-Dioleoyl-sn-glycero-3-phosphocholine
FBS: Fetal Bovine Serum
DMEM: Dulbecco’s Modified Eagle Medium
RPMI-1640: Roswell Park Memorial Institute 1640 Medium
DMSO: Dimethyl Sulfoxide
MTT: 3-(4,5-Dimethylthiazol-2-yl)-2,5-diphenyltetrazolium bromide
PBS: Phosphate-Buffered Saline
IC_50_: Half-Maximal Inhibitory Concentration
LDH: Lactate Dehydrogenase
Rhodamine-PE: Rhodamine-Phosphatidylethanolamine
CMC: Critical Micelle Concentration
DLS: Dynamic Light Scattering
DI: Deionized
H&C NMR: ^1^H and ^13^C Nuclear Magnetic Resonance spectroscopy
SD: Standard Deviation
p-value: Probability Value
ACS: American Chemical Society
Trypsin-EDTA: Trypsin–Ethylenediaminetetraacetic Acid
PPM: Parts Per Million
B: broad
S: singlet
D: doublet
T: triplet
Q: quartet
M: multiplet
δ: Chemical shift
EDCI: 1-Ethyl-3-(3-dimethylaminopropyl) carbodiimide
DMAP: 4-Dimethylaminopyridine
DCM: Dichloromethane
TLC: Thin-Layer Chromatography
CAN: Ceric Ammonium Nitrate
UV: Ultraviolet
R_F_: Retention Factor
ATCC: American Type Culture Collection

