## Supporting Information for "A Saturated Aminotriester Lipid Enhances Amonafide Activity in Human Colorectal Cancer Cells"

CONTENTS

1. General Information SI-2
    a. Materials SI-2
    b. Equipment SI-2

2. Experimental Procedures SI-3
    a. Chemical Synthesis SI-3
    b. Cell Culture SI-19
    c. Cell Viability Assays SI-20
        i. MTT Assay SI-20
        ii. LDH Assay SI-20
    d. Fluorescence-based Cell Uptake Assay SI-21

3. Supplementary Figures SI-22

### 1. General Information

#### a. Materials

Solvents used in all reactions and purification processes were ACS grade or higher and were used without additional purification. They were purchased from AK Scientific, Acros Chemicals, Florida Sunshine Products, or Sigma Aldrich. Deuterated solvents were purchased from Cambridge Isotope Laboratories and were used without further purification. McCoy's 5A Media, Dulbecco's Modified Eagle's Medium (DMEM), RPMI-1640 Medium, penicillin-streptomycin, and 0.25% trypsin-EDTA were all obtained from Tribioscience (Sunnyvale, CA, USA). Fetal bovine serum (FBS) was obtained from Gibco. All other reagents and chemicals were purchased from commercial sources and used without further purification unless otherwise stated.

#### b. Equipment

$^{1}$H and $^{13}$C{$^{1}$H} NMR spectra were acquired on a Varian INOVA 400 MHz nuclear magnetic resonance spectrometer, a Bruker Avance Neo 400 MHz nuclear magnetic resonance spectrometer, or Nanalysis NMReady 60Pro multinuclear benchtop nuclear magnetic resonance spectrometer and were processed on the MestreNova software package. $^{1}$H and $^{13}$C chemical shifts are reported in parts per million (ppm). $^{1}$H chemical shifts are reported relative to the residual solvent peak ($CDCl_3$ = 7.26 ppm) as follows: chemical shift (δ), multiplicity (app = apparent, b = broad, s = singlet, d = doublet, t = triplet, q = quartet, m = multiplet, or combinations thereof), coupling constant(s) in Hz, integration. $^{13}$C chemical shifts are reported relative to the residual solvent peak ($CDCl_3$ = 77.00 ppm). Fluorescence spectra were obtained with the Perkin-Elmer LS-50B Luminescence Spectrophotometer. MTT data was acquired on a Molecular Devices SpectraMax 250 Microplate Spectrophotometer. Cell images were acquired on the Zeiss Axiovert 200 widefield fluorescence microscope with the aperture set to 1/6 and brightfield illumination at 8.2V using two brightfield filters. Dynamic Light Scattering measurements were taken with a Protein Solutions DynaPro-99-E-50 System Dynamic Light Scattering Module.

### 2. Experimental Procedures

#### a. Chemical Synthesis & Characterization

##### Preparation of Stearate Triethanolamine Triester Lipid

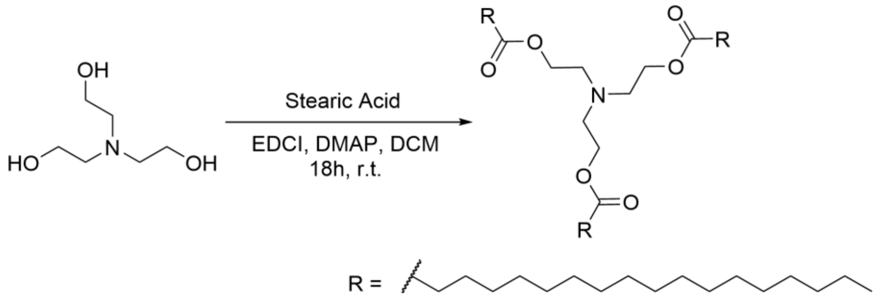


**Chemicals:**
Stearic acid (Acros Chemicals, 98%): used without further purification
Triethanolamine (Florida Sunshine Products, 97%): used without further purification
EDCI (AK Scientific, 98%): used without further purification
DMAP (AK Scientific, >99%): used without further purification
DCM (Stellar Chemical Corp, ACS grade): used without further purification

**Procedure:**
To a 250 mL round bottom flask was added triethanolamine (1.31 g, 8.79 mmol, 1.0 eq) and a Teflon stir bar. The triethanolamine was dissolved in 120mL of DCM, and stearic acid (10.00 g, 35.15 mmol, 4.0 eq), EDCI (6.715 g, 35.15 mmol, 4.0 eq), and DMAP (0.859 g, 7.03 mmol, 0.8 eq) were added. The reaction was allowed to stir at room temperature for 18 hours and was determined complete upon TLC analysis. The reaction was then extracted with one portion of ethyl acetate and KCl brine, where the aqueous layer was then back extracted twice with hexanes, and the three organic layers were combined. The resulting organic layer was dried over anhydrous magnesium sulfate, filtered, and concentrated *in vacuo*. The crude residue was then loaded onto a silica column, where the stearoyl tripod lipid was eluted in a gradient of 3% → 6% ethyl acetate / hexanes to afford the title compound (6.1 g, 73%) as a white wax.

**Characterization Data for Stearate Triethanolamine Triester Lipid**

**TLC:** $R_f$ = 0.60 (10% EtOAc / 90% Hexanes), Non-UV active, periwinkle-blue spot by CAN.

**$^1$H NMR of Stearate Triethanolamine Triester Lipid:** (400 MHz, $CDCl_3$) δ 4.11 (t, *J* = 6.1 Hz, 6H), 2.83 (t, *J* = 6.1 Hz, 6H), 2.33 – 2.26 (m, 6H), 1.61 (d, *J* = 6.8 Hz, 6H), 1.25 (s, 84H), 0.90 – 0.86 (t, *J* = 7.0 Hz, 9H).

**$^{13}$C NMR of Stearate Triethanolamine Triester Lipid:** (101 MHz, $CDCl_3$) δ 173.76, 62.41, 53.31, 34.30, 33.64, 31.94, 29.72, 29.70, 29.68, 29.66, 29.52, 29.38, 29.32, 29.26, 29.21, 24.95, 24.74, 22.71, 14.13, 1.03.

**$^1$H NMR of Stearate Triethanolamine Triester Lipid:** (400 MHz, $CDCl_3$)

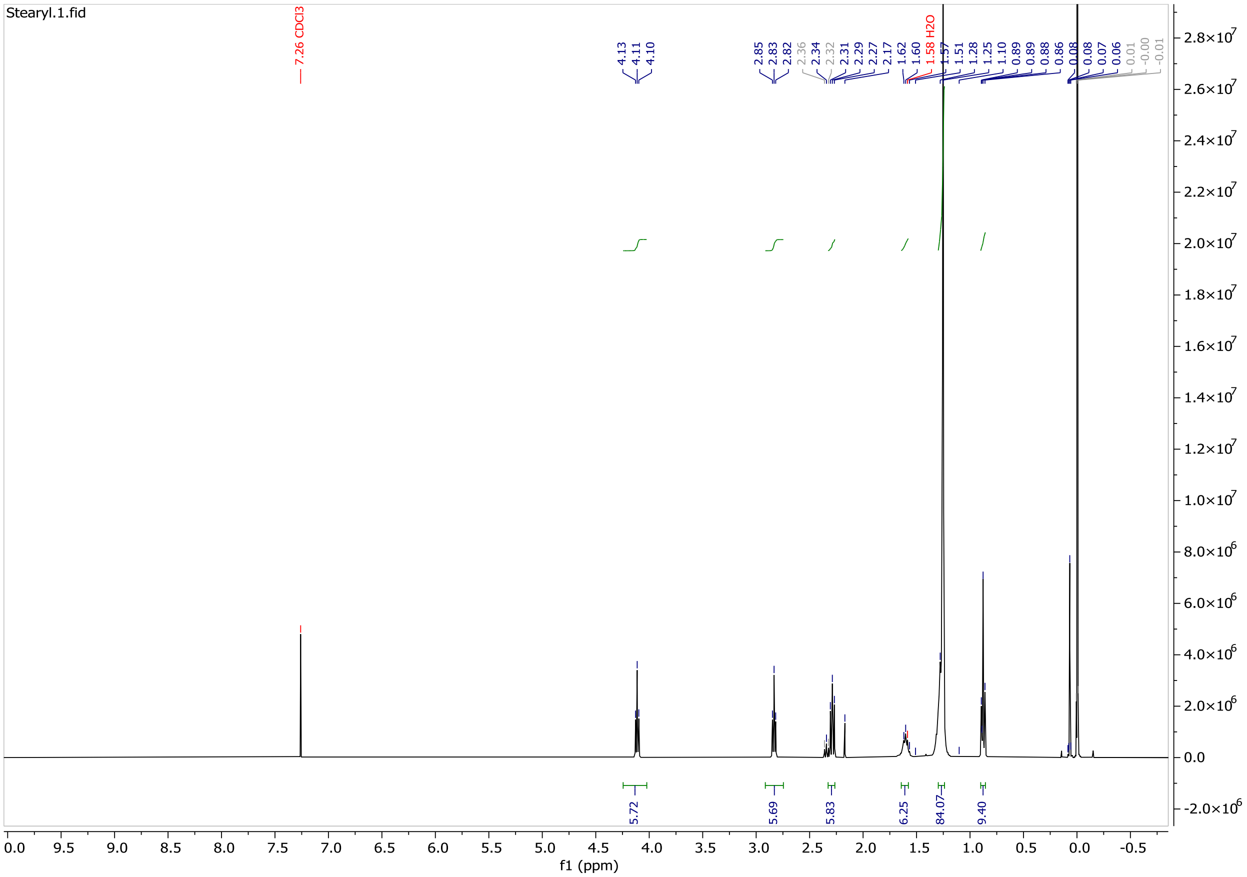

**$^{13}$C NMR of Stearate Triethanolamine Triester Lipid:** (101 MHz, $CDCl_3$)

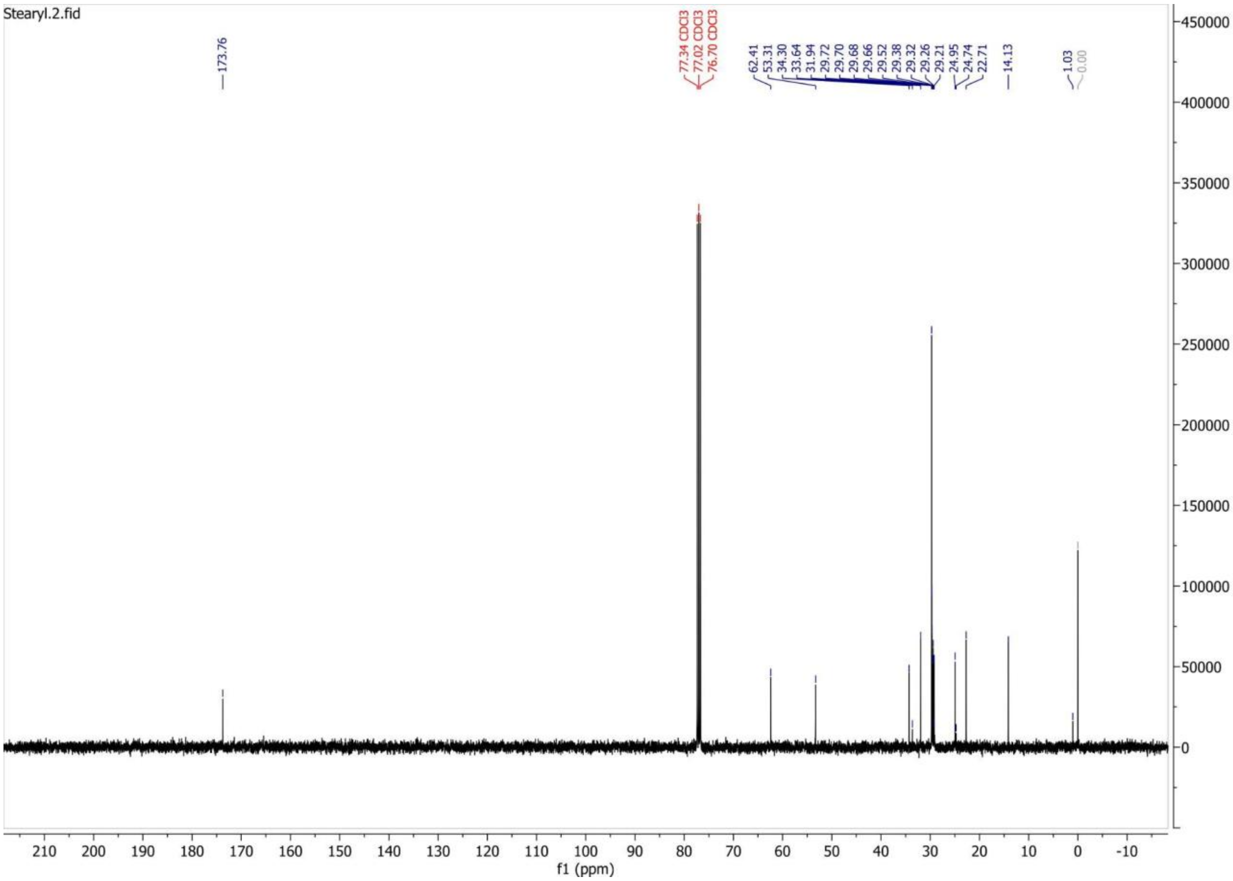

#### Preparation of Oleate Triethanolamine Triester Lipid

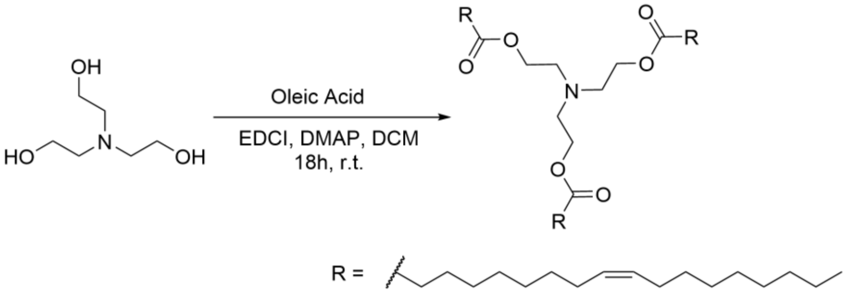


**Chemicals:**

Oleic acid (Acros Chemicals, 98%): used without further purification
Triethanolamine (Sigma Aldrich, 97%): used without further purification
EDCI (AK Scientific, 98%): used without further purification
DMAP (AK Scientific, >99%): used without further purification
DCM (Stellar Chemical Corp, ACS grade): used without further purification

**Procedure:**

To a 250 mL round bottom flask was added triethanolamine (1.32 g, 8.85 mmol, 1.0 eq) and a Teflon stir bar. The triethanolamine was dissolved in 120mL of DCM, and oleic acid (10.00 g, 35.4 mmol, 4.0 eq), EDCI (6.79 g, 35.4 mmol, 4.0 eq), and DMAP (1.08 g, 8.85 mmol, 1.0 eq) were added. The reaction was allowed to stir at room temperature for 18 hours and was determined complete upon TLC analysis. The reaction was then extracted with one portion of ethyl acetate and KCl brine, where the aqueous layer was back extracted with another portion of ethyl acetate, and the two organic layers were combined. The resulting organic layer was dried over anhydrous magnesium sulfate, filtered, and concentrated *in vacuo*. The crude residue was then loaded onto a silica column, where the oleyl tripod lipid was eluted in a gradient of 3% → 10% ethyl acetate / hexanes to afford the title compound (4.32 g, 52% yield) as a yellow oil.

#### Characterization Data for Oleate Triethanolamine Triester Lipid

**TLC:** $R_f$ = 0.50 (10% EtOAc / 90% Hexanes), Non-UV active, denim-blue spot by CAN.

**$^1$H NMR of Oleate Triethanolamine Triester Lipid:** (400 MHz, $CDCl_3$) δ 5.48 – 5.25 (m, 6H), 4.13 (t, J = 6.1 Hz, 6H), 2.85 (t, J = 6.1 Hz, 6H), 2.31 (t, J = 7.6 Hz, 6H), 2.14 – 1.84 (m, 12H), 1.62 (q, J = 7.3 Hz, 6H), 1.36 – 1.16 (m, 60H), 0.92 – 0.87 (m, 9H).

**$^{13}$C NMR of Oleate Triethanolamine Triester Lipid:** (101 MHz, $CDCl_3$) δ 173.70, 130.21, 130.00, 129.73, 127.91, 62.41, 53.32, 34.29, 34.26, 32.62, 31.94, 31.92, 31.80, 31.54, 29.78, 29.72, 29.67, 29.64, 29.63, 29.54, 29.51, 29.38, 29.36, 29.33, 29.31, 29.22, 29.19, 29.16, 29.14, 29.00, 27.23, 27.21, 25.64, 24.95, 24.93, 22.69, 22.67, 22.59, 14.12, 14.08.

**[1]H NMR of Oleate Triethanolamine Triester Lipid:** (400 MHz, $CDCl_3$)

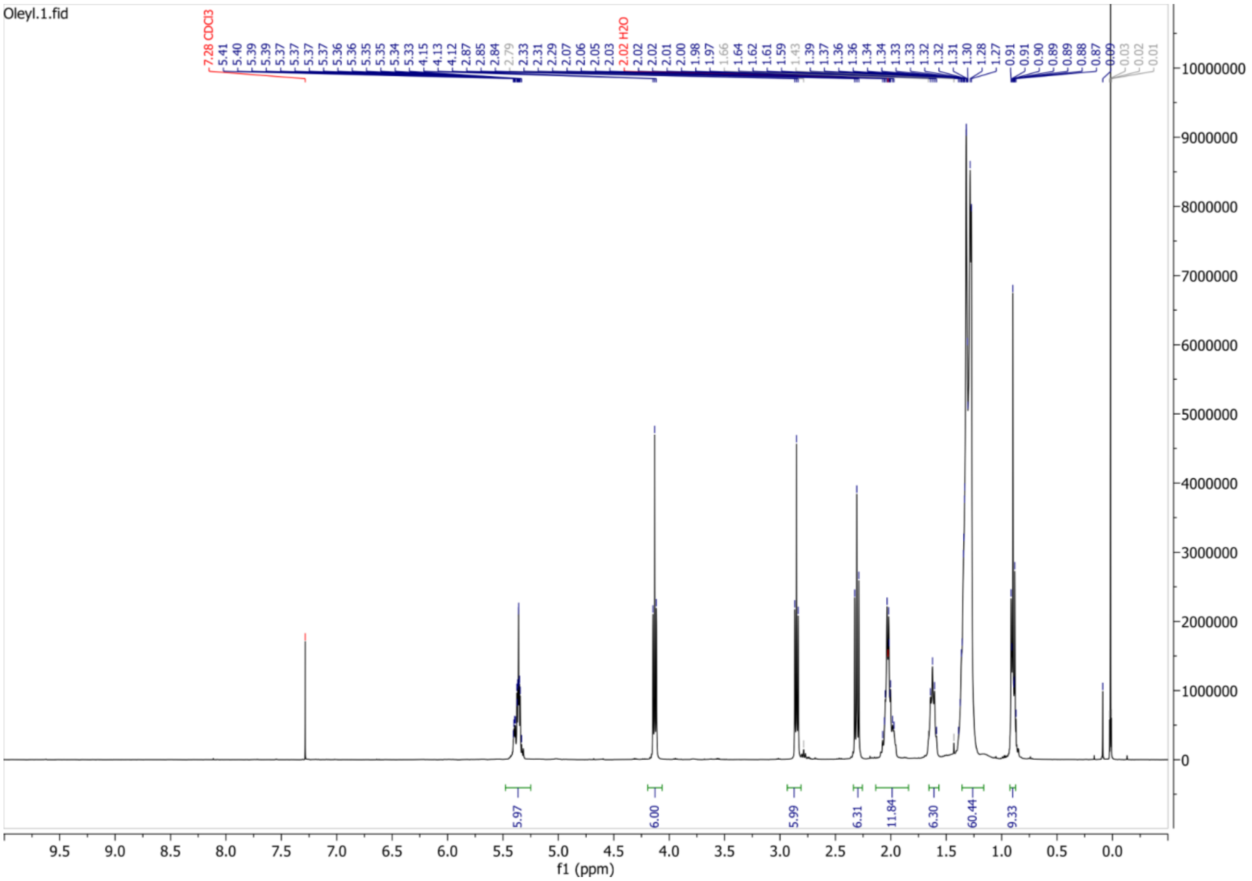

**$^{13}$C NMR of Oleate Triethanolamine Triester Lipid:** (101 MHz, $CDCl_3$)

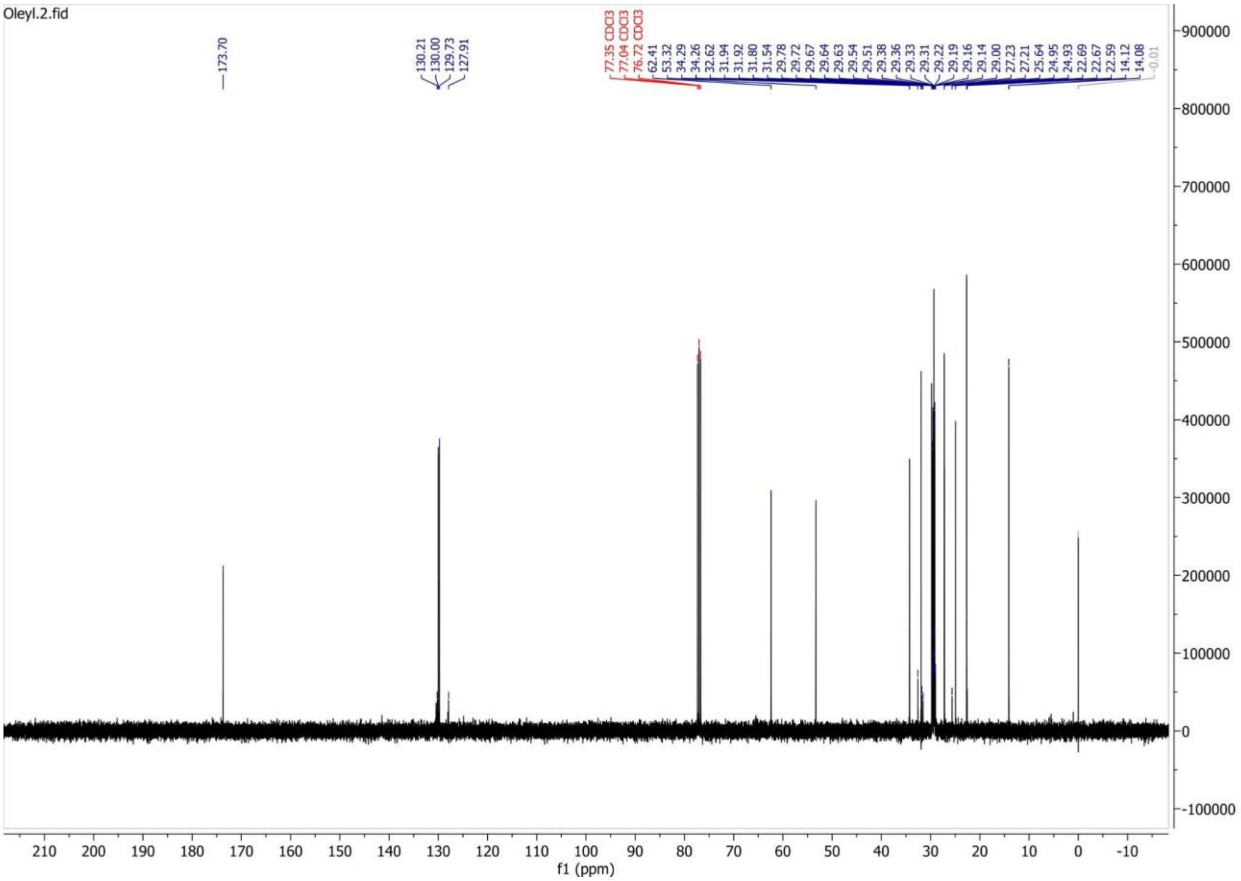

#### Preparation of Linoleate Triethanolamine Triester Lipid

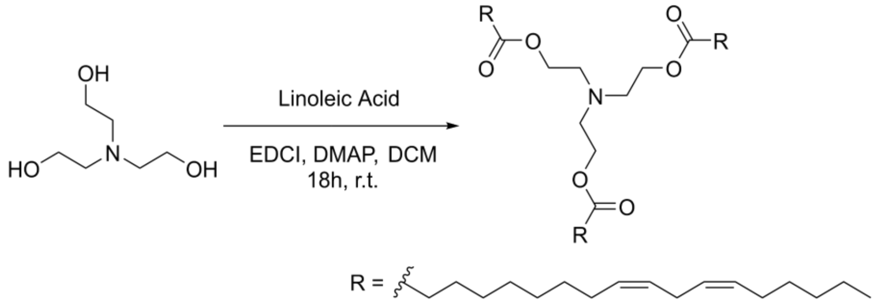


**Chemicals:**
Linoleic acid (AK Scientific, 98%): used without further purification
Triethanolamine (Sigma Aldrich, 97%): used without further purification
EDCI (AK Scientific, 98%): used without further purification
DMAP (AK Scientific, >99%): used without further purification
DCM (Stellar Chemical Corp, ACS grade): used without further purification

**Procedure:**
To a 250mL round bottom flask was added triethanolamine (1.329 g, 8.91 mmol, 1.0 eq) and a Teflon stir bar. The triethanolamine was dissolved in 120 mL of DCM, and linoleic acid (10.00 g, 35.65 mmol, 4.0 eq), EDCI (6.81 g, 35.65 mmol, 4.0 eq), and DMAP (868.9 mg, 7.130 mmol, 0.8 eq) were added. The reaction was allowed to stir at room temperature for 18 hours and was determined complete upon TLC analysis. The reaction was then extracted with one portion of ethyl acetate and 1 M sodium bicarbonate, where the aqueous layer was then back extracted twice with ethyl acetate. The resulting organic layer was dried over anhydrous sodium sulfate, filtered, and concentrated *in vacuo*. The crude residue was then loaded onto a silica column, where the linoleyl tripod lipid was eluted in a gradient of 3% → 6% ethyl acetate / hexanes to afford the title compound (6.1g, 73%) as a white waxy solid.

**Characterization Data for Linoleate Triethanolamine Triester Lipid**

**TLC:** $R_f$ = 0.60 (10% EtOAc / 90% Hexanes), Non-UV active, periwinkle-blue spot by CAN.

**$^{1}$H NMR of Linoleate Triethanolamine Triester Lipid:** (400 MHz, $CDCl_3$) δ 5.46 – 5.21 (m, 6H), 4.10 (t, *J* = 6.1 Hz, 6H), 2.82 (t, *J* = 6.1 Hz, 6H), 2.28 (t, *J* = 7.6 Hz, 6H), 2.02 (dd, *J* = 15.2, 7.1 Hz, 6H), 1.60 (p, *J* = 7.4 Hz, 6H), 1.43 – 1.10 (m, 60H), 0.87 (td, *J* = 6.9, 4.0 Hz, 9H).

**$^{13}$C NMR of Linoleate Triethanolamine Triester Lipid:** (101 MHz, $CDCl_3$) δ 173.71, 173.68, 173.67, 130.20, 130.00, 129.99, 129.72, 128.05, 127.91, 62.40, 53.32, 34.28, 34.25, 31.94, 31.53, 29.78, 29.72, 29.70, 29.67, 29.64, 29.62, 29.54, 29.50, 29.38, 29.36, 29.33, 29.31, 29.21, 29.19, 29.15, 29.14, 27.23, 27.20, 27.18, 25.63, 24.94, 24.92, 22.70, 22.69, 22.58, 14.12.

**[1]H NMR of Linoleate Triethanolamine Triester Lipid:** (400 MHz, $CDCl_3$)

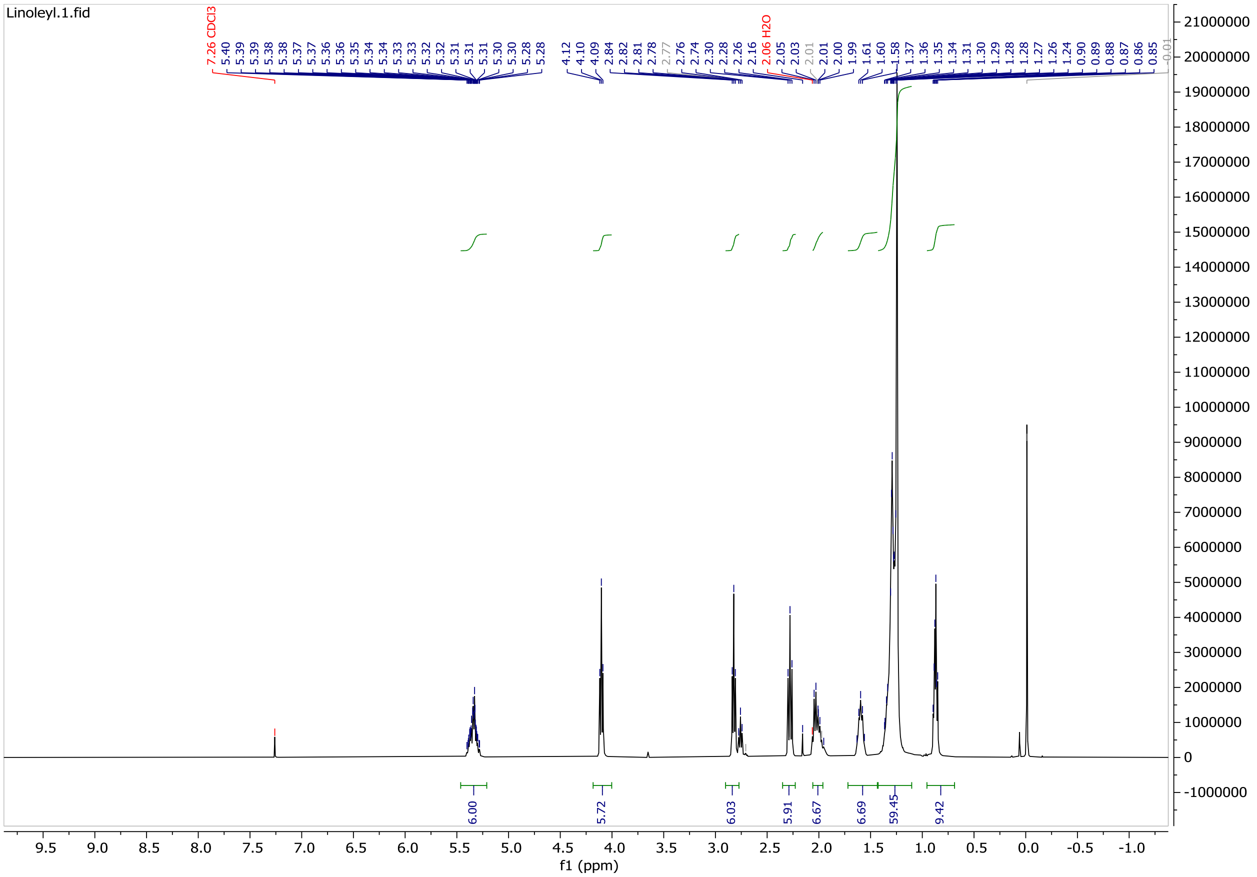

**$^{13}$C NMR of Linoleate Triethanolamine Triester Lipid:** (101 MHz, $CDCl_3$)

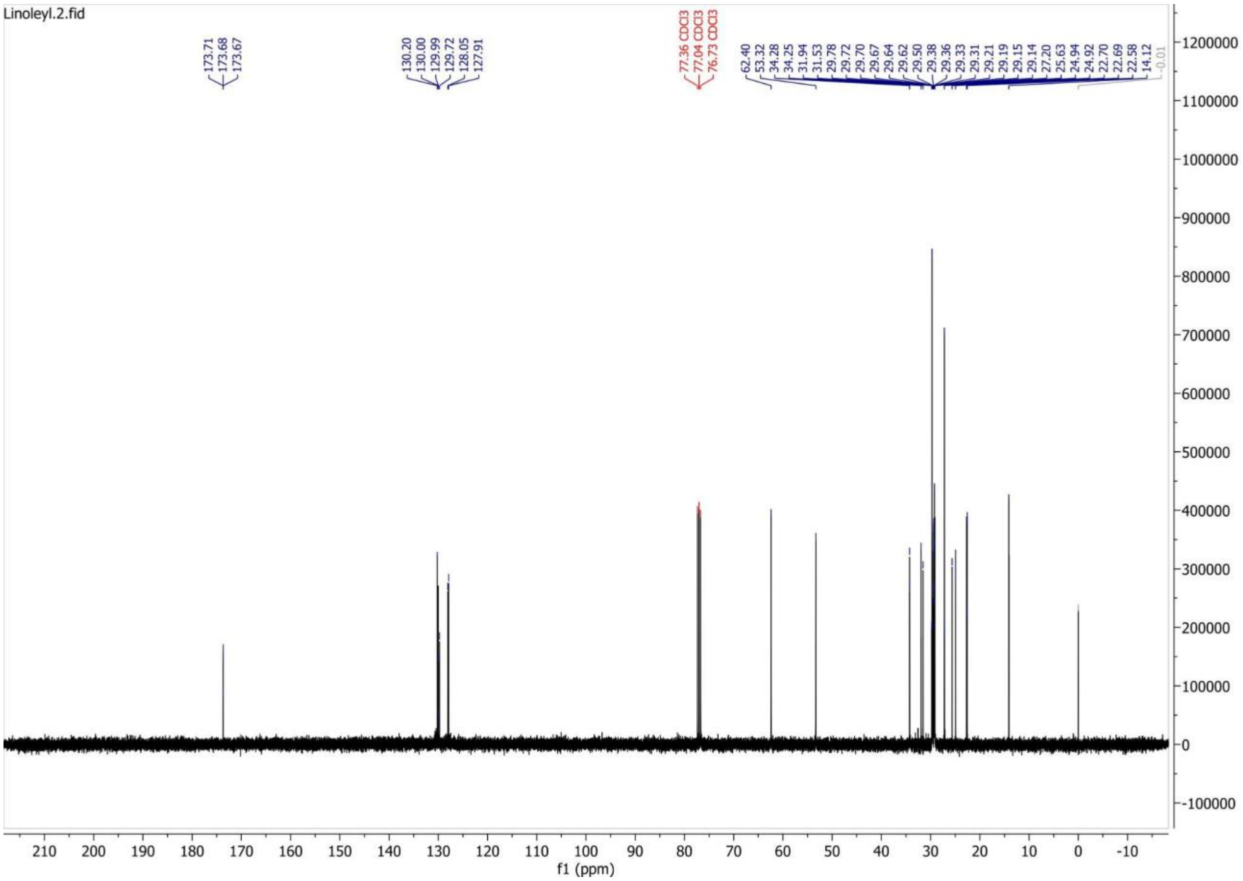

#### Preparation of Elaidate Triethanolamine Triester Lipid

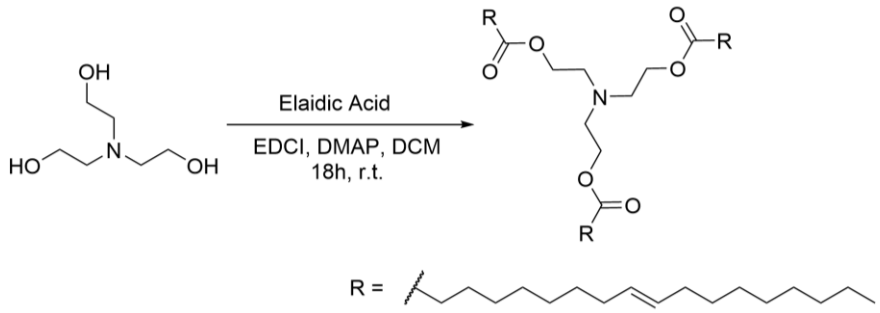


**Chemicals:**
Elaidic acid (Acros Chemicals, 98%): used without further purification
Triethanolamine (Sigma Aldrich, 97%): used without further purification
EDCI (AK Scientific, 98%): used without further purification
DMAP (AK Scientific, 99%): used without further purification
DCM (Stellar Chemical Corp, ACS grade): used without further purification

**Procedure:**
To a 250 mL round bottom flask was added triethanolamine (1.32 g, 8.85 mmol, 1.0 eq) and a Teflon stir bar. The triethanolamine was dissolved in 120 mL of DCM, and elaidic acid (10.00 g, 35.40 mmol, 4.0 eq), EDCI (6.79 g, 35.40 mmol, 4.0 eq), and DMAP (864.96 mg, 7.08 mmol, 0.8 eq) were added. The reaction was allowed to stir at room temperature for 18 hours and was determined complete upon TLC analysis. The reaction was then extracted with one portion of ethyl acetate and KCl brine, where the aqueous layer was then back extracted twice with hexanes. The resulting organic layer was dried over anhydrous magnesium sulfate, filtered, and concentrated *in vacuo*. The crude residue was then loaded onto a silica column, where the elaidyl tripod lipid was eluted in a gradient of 3% → 6% ethyl acetate / hexanes to afford the title compound (6.1g, 73%) as a white wax.

**Characterization Data for Elaidate Triethanolamine Triester Lipid**

**TLC:** $R_f$ = 0.60 (10% EtOAc / 90% Hexanes), Non-UV active, periwinkle-blue spot by CAN.

**$^1$H NMR of Elaidate Triethanolamine Triester Lipid:** (400 MHz, $CDCl_3$) δ 5.41 – 5.32 (m, 6H), 4.11 (t, *J* = 6.1 Hz, 6H), 2.83 (t, *J* = 6.1 Hz, 6H), 2.28 (t, *J* = 7.6 Hz, 6H), 1.95 (q, *J* = 6.3 Hz, 12H), 1.60 (p, *J* = 7.3 Hz, 6H), 1.35 – 1.24 (m, 60H), 0.90 – 0.85 (t, *J* = 7.0 Hz, 9H).

**$^{13}$C NMR of Elaidate Triethanolamine Triester Lipid:** (101 MHz, $CDCl_3$) δ 173.72, 130.47, 130.20, 62.41, 53.32, 34.27, 32.62, 32.58, 31.92, 29.67, 29.61, 29.51, 29.33, 29.21, 29.16, 29.00, 24.94, 22.69, 14.13.

**$^{1}$H NMR of Elaidate Triethanolamine Triester Lipid:** (400 MHz, $CDCl_3$)

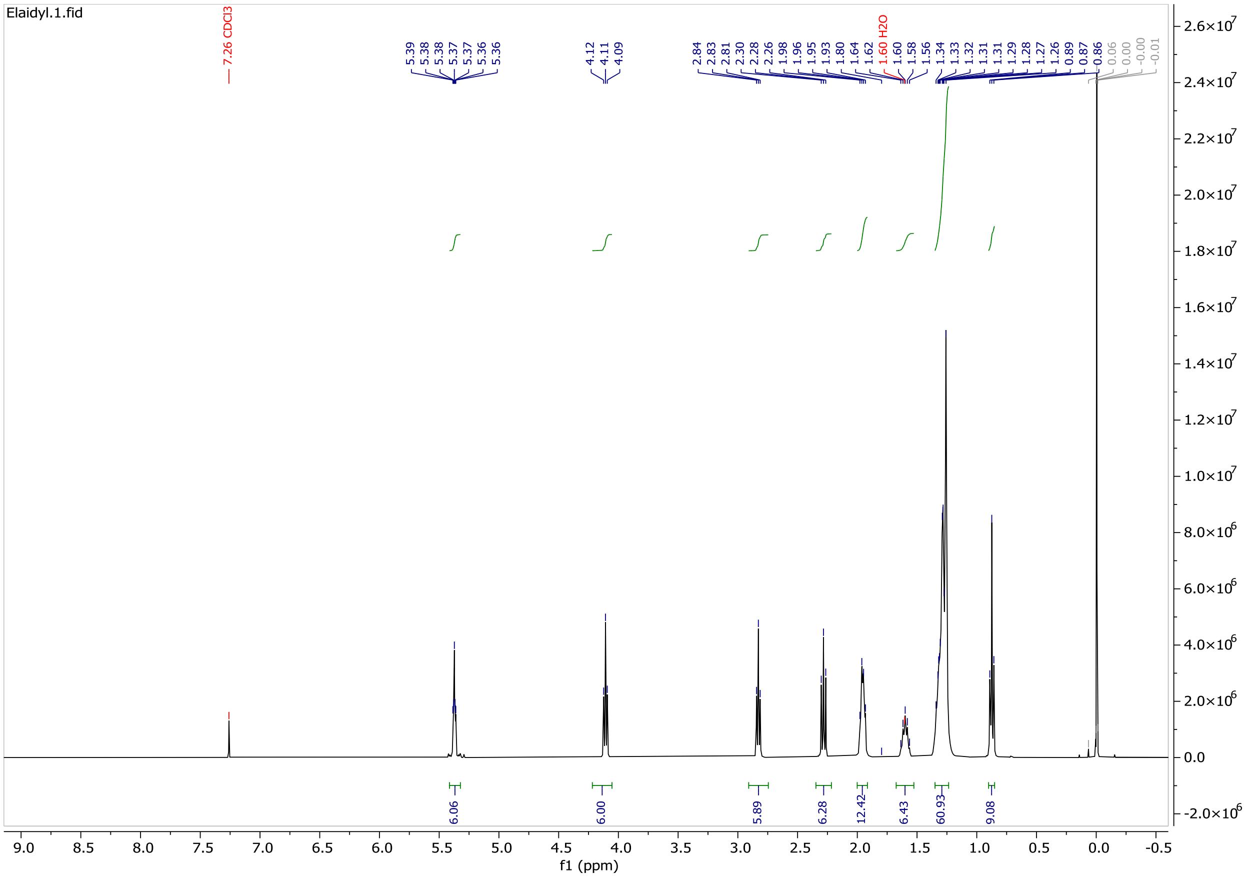

**$^{13}$C NMR of Elaidate Triethanolamine Triester Lipid:** (101 MHz, $CDCl_3$)

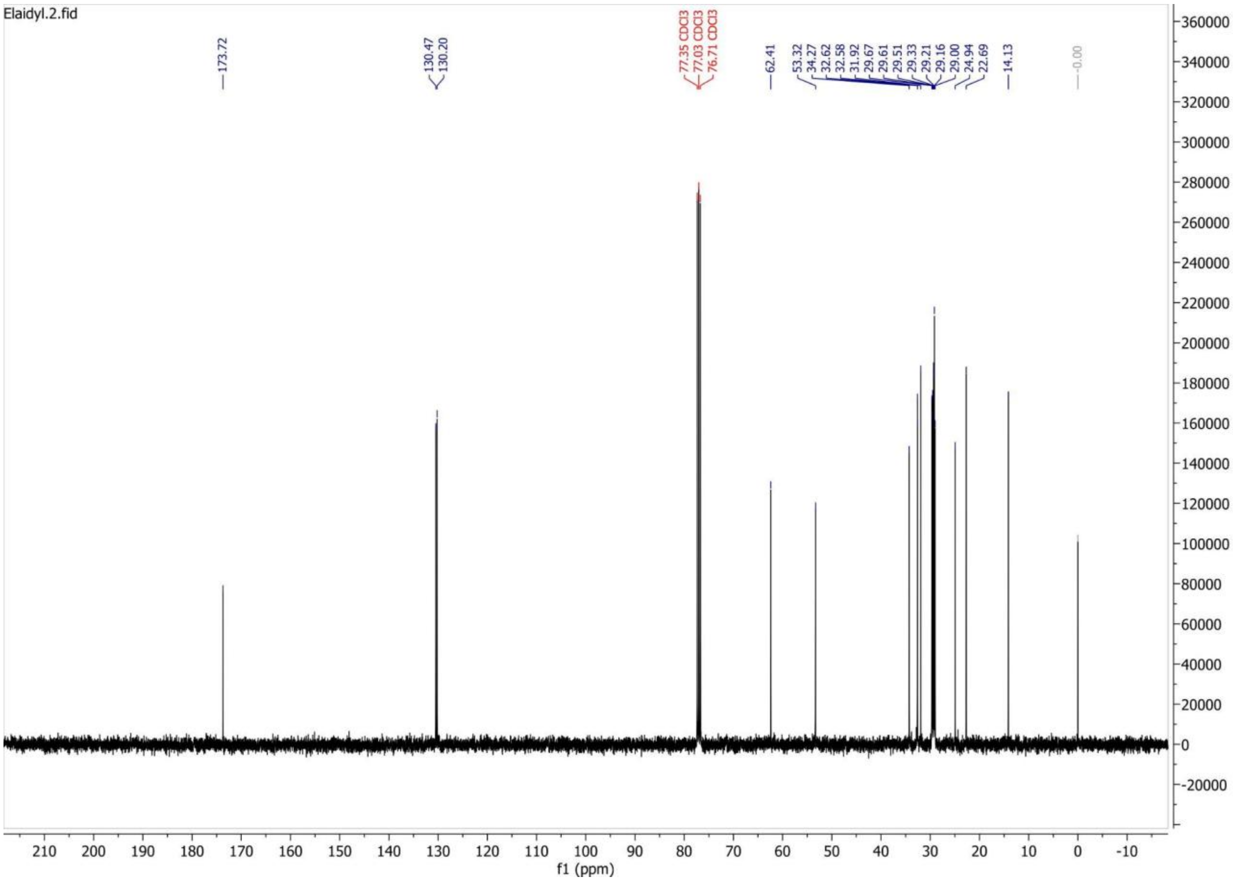

#### b. Cell Culture

HCT-116 and HT-29 human colorectal cancer cell lines were obtained from the European Collection of Authenticated Cell Cultures (ECACC) and cultured in McCoy's 5A Medium (Tribioscience) supplemented with 10% v/v fetal bovine serum (FBS, Gibco) and 1% v/v 100x penicillin-streptomycin (Tribioscience). CT26, a murine colorectal carcinoma cell line, was obtained from American Type Culture Collection (ATCC), and cultured in RPMI-1640 Medium (Tribioscience) supplemented with 10% v/v
fetal bovine serum (FBS, Gibco) and 1% v/v 100x penicillin-streptomycin (Tribioscience). A549, a human non-small cell lung carcinoma cell line, was maintained in High Glucose Dulbecco's Modified Eagle Medium (DMEM) (Hygia Reagents) supplemented with 10% fetal bovine serum (Gibco) and 1% penicillin-streptomycin (Tribioscience). HCT-116, HT-29, CT26, and A549 cells were cultured in T25 and T75 flasks (Corning). Cultures were kept at 37°C in a humidified incubator (5.0% $CO_2$) and maintained by splitting cells at a ratio of 1:3 to 1:6 based on confluence and cultured up to passage 20.

#### c. Cell Viability

##### i. *MTT Assay*

Cell viability was determined by 3-(4,5-dimethylthiazol-2-yl)-2,5-diphenyltetrazolium bromide (MTT) assays through literature reported protocols. HCT-116 and HT-29 cells were seeded at 70% confluency in McCoy's 5A Medium and supplemented with 10% v/v fetal bovine serum (FBS, Gibco) and 1% v/v 100x penicillin-streptomycin (Tribioscience). A549 and CT26 cells were seeded at 70% confluency in High Glucose Dulbecco's Modified Eagle Medium (DMEM) and RPMI-1640 respectively, supplemented with 10% v/v fetal bovine serum (FBS, Gibco) and 1% v/v 100x penicillin-streptomycin (Tribioscience). Following the 24-hour incubation period, a drug medium solution was prepared by pre-formulating our triethanolamine triester lipids with select chemotherapeutics. Equivalent volumes of drug and 50mM lipid solutions were mixed in a v-bottom 96-well plate, which were added to media in a deep-well plate. 100 µL of the drugged media was then added to each well to reach desired drug concentrations of 25 µM, 5 µM, 2.5 µM, 500 nM, 250 nM, 50 nM, 25 nM, and 5 nM (0.5% v/v DMSO). A negative control of 0.5% v/v DMSO was also included. The plates were then incubated at 37°C (5.0% $CO_2$) for 96 hours, after which 10 µL of a fresh solution of 3-(4,5-dimethylthiazol-2-yl)-2,5-diphenyltetrazolium bromide (MTT) (AK Scientific) in 1x phosphate-buffered saline (PBS) (5 mg/mL) was added to all wells. After mixing, the plates were allowed to incubate for 1-3 hours. The cell media was then aspirated, and 100 µL of DMSO was added to each well and mixed until all formazan crystals were fully solubilized. Absorbance was measured with a Molecular Devices SpectraMax® Plus 384 Microplate Reader at 490 nm. Cell viability was calculated and normalized against the negative control, and $IC_{50}$ values were determined using GraphPad Prism 10.4.1 with an inhibition regression analysis.

##### ii. *Lactate Dehydrogenase (LDH) Assay*

Extracellular LDH activity, released due to a loss of cell membrane integrity, was measured using a previously reported Cold Spring Harbor Protocol. HT-29 cells were seeded at 70% confluency in McCoy's 5A Medium and supplemented with 10% v/v fetal bovine serum (FBS, Gibco) and 1% v/v 100x penicillin-streptomycin (Tribioscience). A549 and CT26 cells were seeded at 70% confluency in High Glucose Dulbecco's Modified Eagle Medium (DMEM) and RPMI-1640 respectively, supplemented with 10% v/v fetal bovine serum (FBS, Gibco) and 1% v/v 100x penicillin-streptomycin (Tribioscience). Following the 24-hour incubation period, a drug medium solution was prepared by pre-formulating our triethanolamine triester lipids with select chemotherapeutics. Equivalent volumes of drug and lipid solutions were mixed in a v-bottom 96-well plate, which were added to media in a deep-well plate. 100 µL of the drugged media was then added to each well to reach desired concentrations of 25 µM, 5 µM, 2.5 µM, 500 nM, 250 nM, 50 nM, 25 nM, and 5 nM (0.5% v/v DMSO). A negative control of 0.5% v/v DMSO was also included. The plates were then incubated at 37°C (5.0% $CO_2$) for 96 hours. At 96 hours, to a negative control of cells drugged with DMSO was added 10 µL of lysis solution (9% v/v Triton X-100) for 5 minutes to act as a positive control for maximum LDH release. 50 µL of supernatant cell media from each treatment was then moved to a non-tissue culture treated 96-well plate and allowed to react with 50 µL of the LDH substrate solution (L-(+)-lactic acid (0.054 M), β-NAD+ (1.30 mM), 1-methoxy-5-methylphenazinium methyl sulfate (0.28 mM), and 2-p-iodophenyl-3-p-nitrophenyl tetrazolium chloride (INT) solution (0.66 mM) dissolved in 0.2 M tris-HCl buffer (pH 8.2)), for 45 min at 37°C (5.0%

$CO_2$) protected from light. Absorbance was measured with a Molecular Devices SpectraMax® Plus 384 Microplate Reader at 490 nm.

#### d. Fluorescence-based Cell Uptake Assay

##### Procedure 1

Cell cultures were prepared, referencing the protocol above. Cells were seeded either in a 96-well flat bottom tissue culture treated plate (Thermo Scientific) or an 8-well imaging plate and incubated for 24 hours at 37°C in a humidified incubator (5 % $CO_2$). Following the 24-hour incubation period, a drug medium solution was prepared by adding DMSO solutions of Amonafide co-delivered with our triethanolamine triester lipids or CaP-Lipid complexes to reach a concentration of 500 µM. 1 µL of the lipid-drug solution was added to 100 µL of cell-specific culture media per well to reach a final concentration of 5 µM. The cells were subsequently incubated at 37°C for 24-hours before the media was aspirated off and replaced with 50 µL 1x PBS. Images were collected with the Zeiss Axiovert 200 widefield fluorescence microscope with the aperture set to ⅙ and brightfield illumination at 4.25V using two brightfield filters with a 32x lens objective (Objective LD A-Plan 32x/0.4 Ph1, item no: 441251-9910-000).

##### Procedure 2

Human colorectal carcinoma (HCT-116) and murine colorectal carcinoma (CT26) cell lines were cultured and utilized to evaluate the fluorescence localization of amonafide in the presence of four lipid formulations. Stock solutions consisted of 10 mM amonafide dissolved in dimethyl sulfoxide (DMSO), four lipid formulations, tris-stearate, tris-oleate, tris-linoleate, tris-elaidate prepared in phosphate-buffered saline (PBS), and 1 mg/mL rhodamine. For each experimental condition, 10 µL of amonafide solution and 20 µL of the designated lipid formulation were added to individual wells. Experiments were conducted three times for each lipid formulation and cell type. Each trial consisted of four treatment groups: (1) amonafide alone, (2) amonafide followed by lipid addition after a 5 min period, (3) lipid alone, and (4) lipid followed by amonafide addition after a 5 min period. Thus, each lipid formulation was evaluated using 12 wells (4 treatment groups 3 replicates). Across all four lipid formulations, a total of 48 wells were analyzed per cell line, resulting in 96 experimental wells across both HCT-116 and CT26 cell types. Immediately prior to imaging, approximately 150 µL of medium was carefully removed from each well, leaving a minimal volume sufficient to prevent cellular desiccation during image acquisition. Wells were then promptly transferred for microscopy. Fluorescence microscopy was performed using a 10x objective lens. For each well, three images were acquired from the same field of view. First, a brightfield image was captured using the white filter taken using halogen. Then, fluorescence images are taken using the green filter set followed by the red filter set, ensuring that all images were collected from the identical imaging location. Fluorescence was used exclusively during the green and red channel images. A total of three images (brightfield, green fluorescence, and red fluorescence) were taken for every experimental well and used for analysis.

### 3. Supplementary Figures

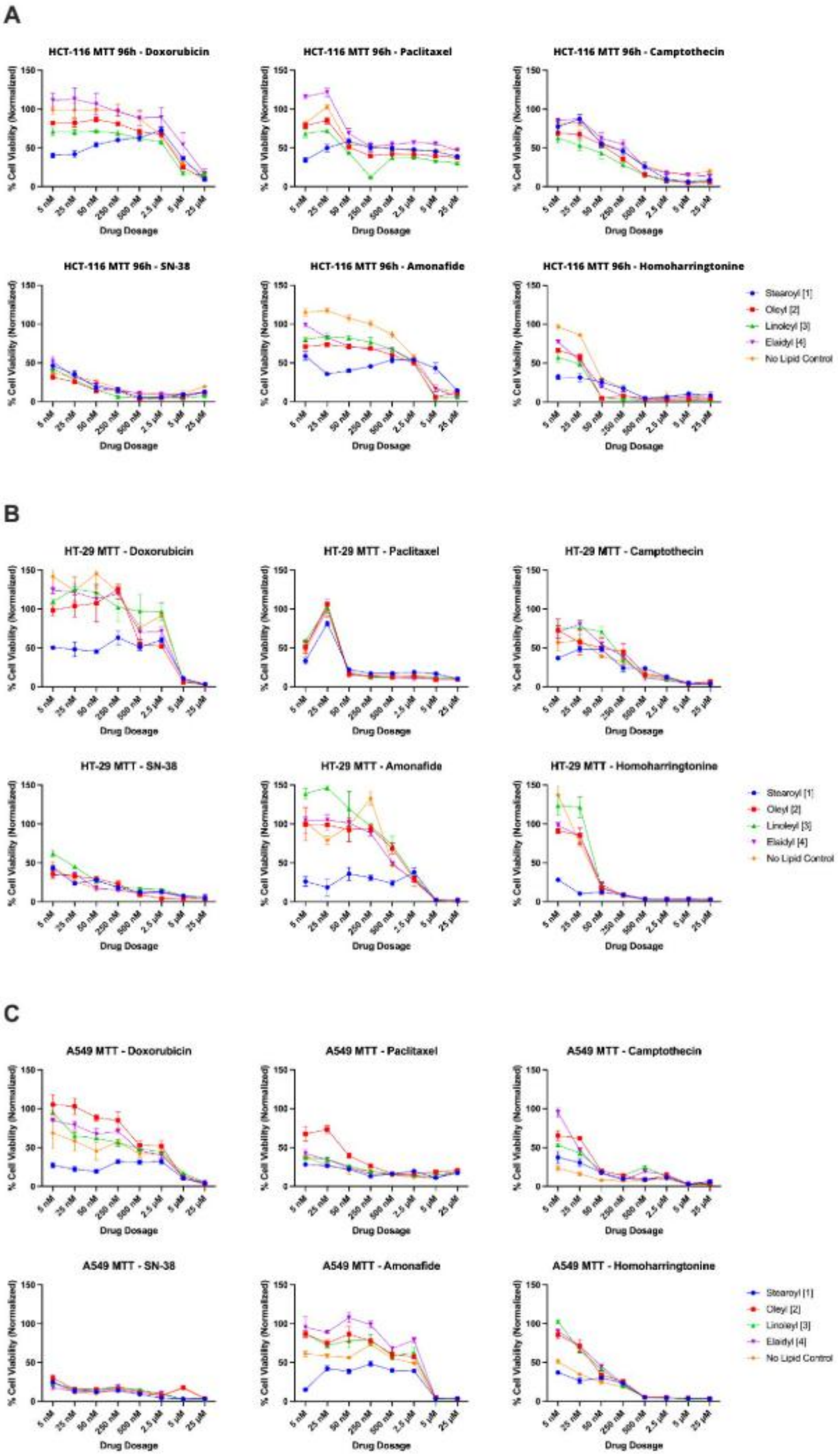

A
HCT-116 MTT 96h - Doxorubicin
HCT-116 MTT 96h - Paclitaxel
HCT-116 MTT 96h - Camptothecin
HCT-116 MTT 96h - SN-38
HCT-116 MTT 96h - Amonafide
HCT-116 MTT 96h - Homoharringtonine
B
HT-29 MTT - Doxorubicin
HT-29 MTT - Paclitaxel
HT-29 MTT - Camptothecin
HT-29 MTT - SN-38
HT-29 MTT - Amonafide
HT-29 MTT - Homoharringtonine
C
A549 MTT - Doxorubicin
A549 MTT - Paclitaxel
A549 MTT - Camptothecin
A549 MTT - SN-38
A549 MTT - Amonafide
A549 MTT - Homoharringtonine
% Cell Viability (Normalized)
0
50
100
150
5 nM
25 nM
50 nM
250 nM
500 nM
2.5 µM
5 µM
25 µM
Drug Dosage
Stearoyl [1]
Oleyl [2]
Linoleyl [3]
Elaidyl [4]
No Lipid Control

**SI 1.** Anti-proliferative in vitro evaluation of doxorubicin, paclitaxel, camptothecin, SN-38, amonafide, and homoharringtonine when co-delivered with *tris*-stearate, *tris*-oleate, *tris*-linoleate, *tris*-elaidate, and a no lipid control in HCT-116, HT-29, and A549 mammalian cell lines.

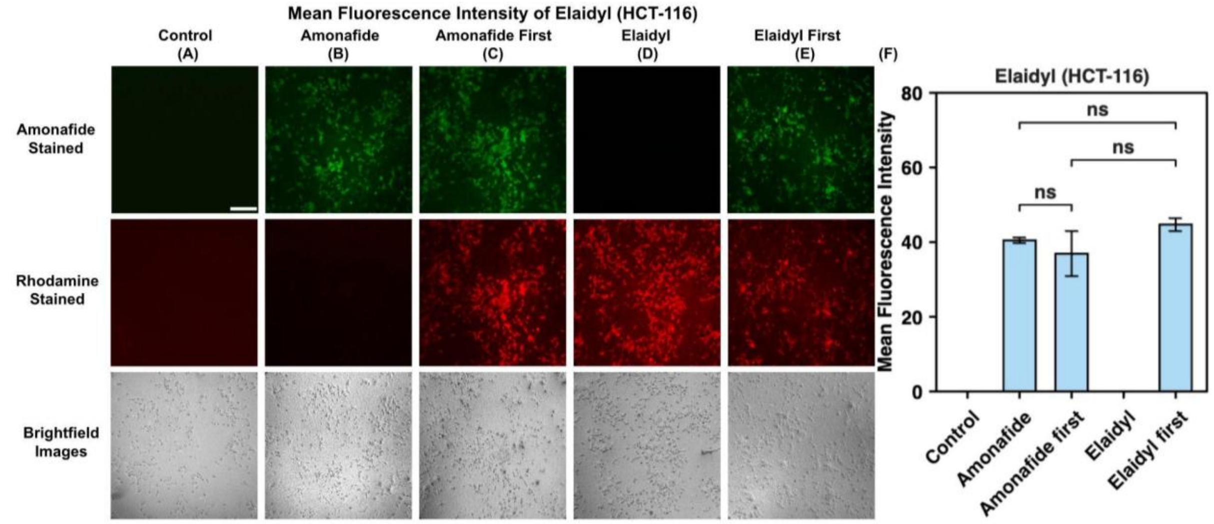


**SI 2.** Fluorescent microscopy images and mean fluorescence intensity graphs of amonafide and *tris*-elaidate in the HCT-116 cell line. The green channel denotes the location of amonafide intracellular delivery, the red channel indicates the presence of rhodamine-tagged *tris*-elaidate, and the brightfield is included to verify the location of the fluorescent-dyed lipid cells. Scale bars are 50 µm **(A)**.

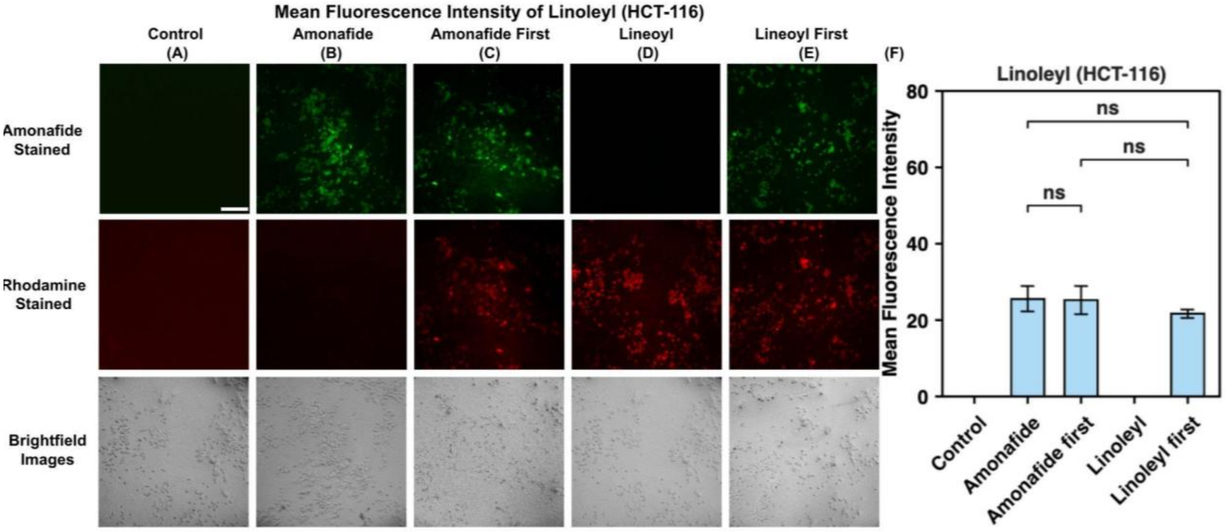


**SI 3.** Fluorescent microscopy images and mean fluorescence intensity graphs of amonafide and *tris*-linoleate in the HCT-116 cell line. The green channel denotes the location of amonafide intracellular delivery, the red channel indicates the presence of rhodamine-tagged *tris*-lineolate, and the brightfield is included to verify the location of the fluorescent-dyed lipid cells. Scale bars are 50 µm **(A)**.

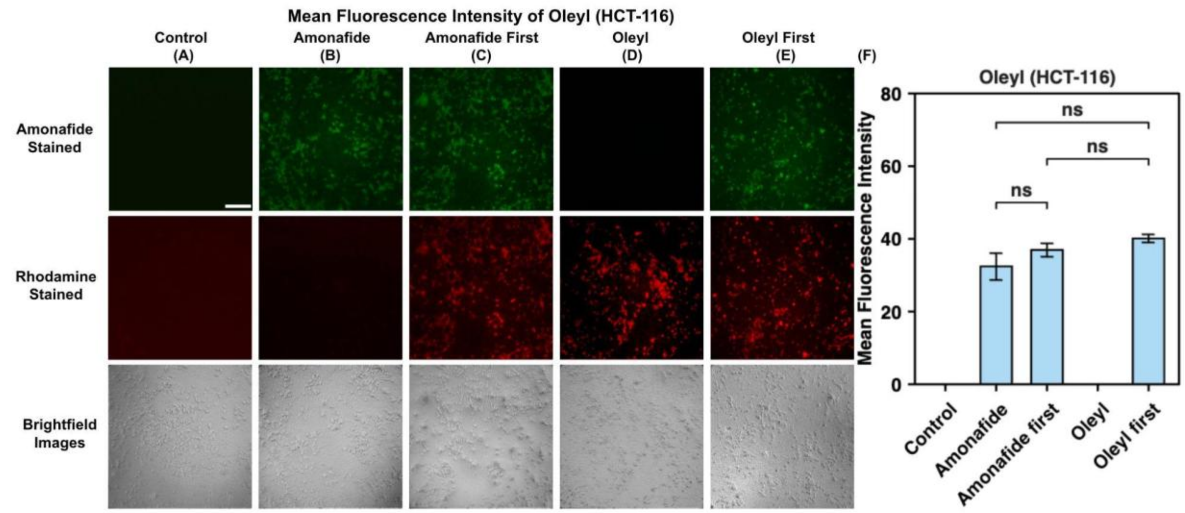


**SI 4.** Fluorescent microscopy images and mean fluorescence intensity graphs of amonafide and *tris*-oleate in the HCT-116 cell line. The green channel denotes the location of amonafide intracellular delivery, the red channel indicates the presence of rhodamine-tagged *tris*-oleate, and the brightfield is included to verify the location of the fluorescent-dyed lipid cells. Scale bars are 50 µm **(A)**.

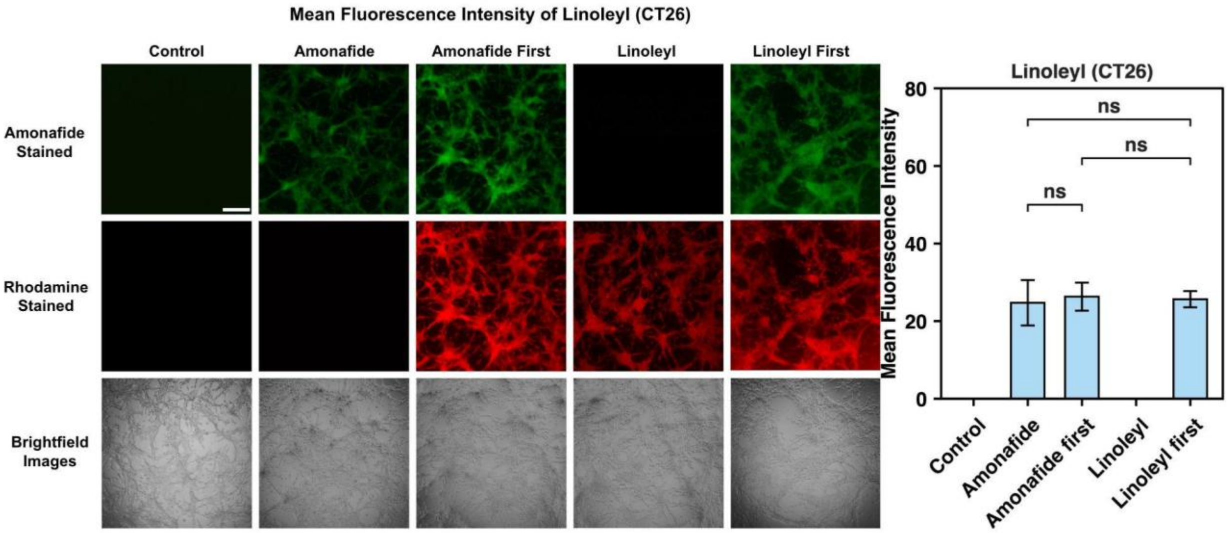


**SI 5.** Fluorescent microscopy images and mean fluorescence intensity graphs of amonafide and *tris*-linoleate in the CT26 cell line. The green channel denotes the location of amonafide intracellular delivery, the red channel indicates the presence of rhodamine-tagged *tris*-linoleate, and the brightfield is included to verify the location of the fluorescent-dyed lipid cells. Scale bars are 50 µm **(A)**.

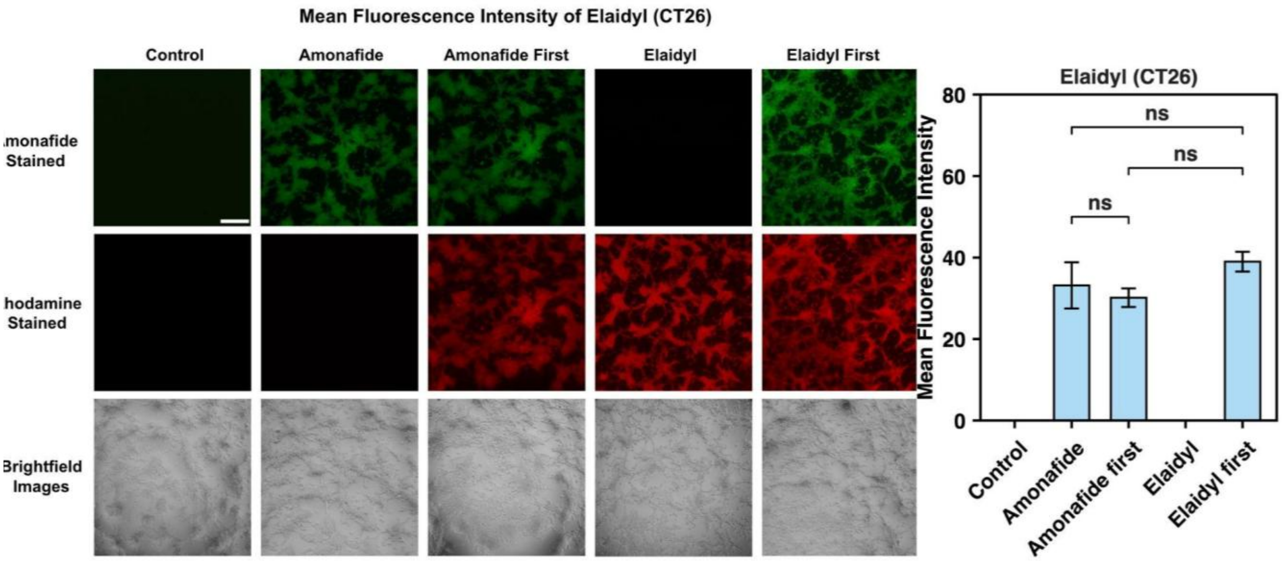


**SI 6.** Fluorescent microscopy images and mean fluorescence intensity graphs of amonafide and *tris*-elaidate in the CT26 cell line. The green channel denotes the location of amonafide intracellular delivery, the red channel indicates the presence of rhodamine-tagged *tris*-elaidate, and the brightfield is included to verify the location of the fluorescent-dyed lipid cells. Scale bars are 50 µm **(A)**.

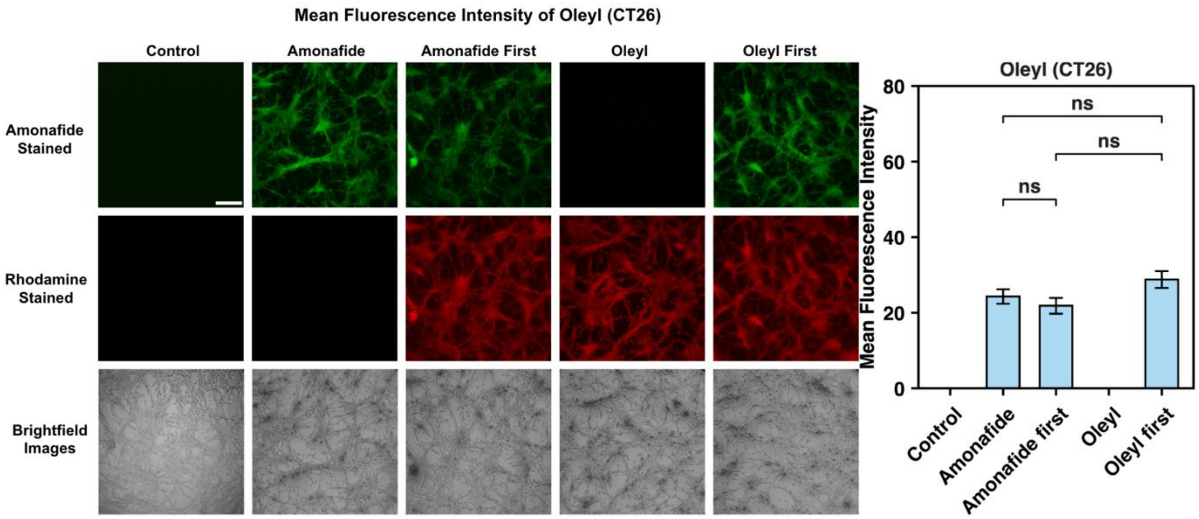


**SI 7.** Fluorescent microscopy images and mean fluorescence intensity graphs of amonafide and oleyl in the CT26 cell line. The green channel denotes the location of amonafide intracellular delivery, the red channel indicates the presence of rhodamine-tagged *tris*-oleate, and the brightfield is included to verify the location of the fluorescent-dyed lipid cells. Scale bars are 50 µm **(A)**.

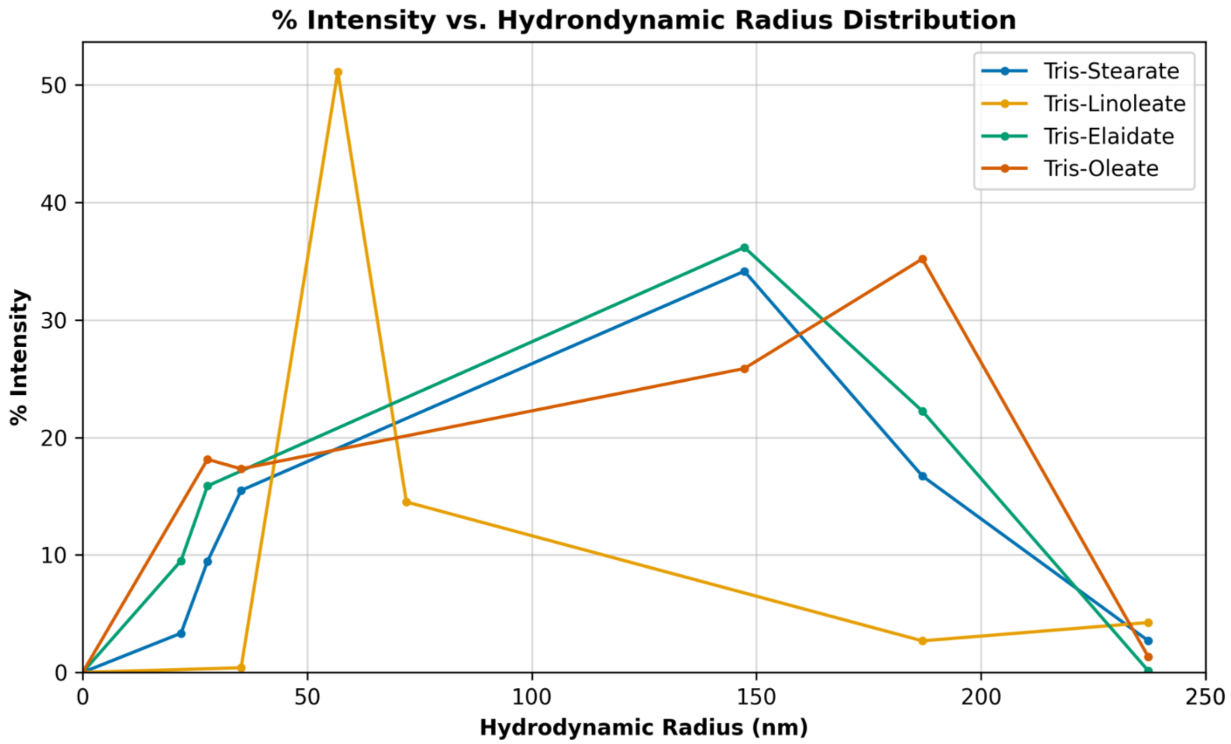


**SI 8.** Percent Intensity against Hydrodynamic Radius for four aminotriester lipids without the presence of amonafide.
